# Application of high-throughput, high-depth, targeted single-nucleus DNA sequencing in pancreatic cancer

**DOI:** 10.1101/2022.03.06.483206

**Authors:** Haochen Zhang, Elias-Ramzey Karnoub, Shigeaki Umeda, Ronan Chaligné, Ignas Masilionis, Caitlin A. McIntyre, Akimasa Hayashi, Palash Sashittal, Amanda Zucker, Katelyn Mullen, Alvin Makohon-Moore, Christine A. Iacobuzio-Donahue

## Abstract

Despite insights gained by bulk DNA sequencing of cancer it remains challenging to resolve the admixture of normal and tumor cells, and/or of distinct tumor subclones; high throughput single-cell DNA sequencing circumvents these and brings cancer genomic studies to higher resolution. However, its application has been limited to liquid tumors or a small batch of solid tumors, mainly because of the lack of a scalable workflow to process solid tumor samples. Here we optimized a highly automated nuclei extraction workflow that achieved fast and reliable targeted single-nucleus DNA library preparation of 38 samples from 16 pancreatic adenocarcinoma (PDAC) patients, with an average library yield per sample of 2867 single nuclei. We demonstrate that this workflow not only performs well using low cellularity or low tumor purity samples but reveals novel genomic evolution patterns of PDAC as well.

## Introductionn

The field of single-cell genomics, since its advent about 10 years ago^1,2^, has been striving to increase the throughput and resolution of cancer research. Single-cell DNA sequencing (scDNA-seq) offers many advantages over traditional “bulk” DNA-seq. Most importantly, it circumvents the issue of “mixed signals”^3^, i.e. the admixture of normal and tumor cells, and/or of distinct tumor subclones. Solving the former allows for much higher sensitivity in calling rare genetic events, which opens opportunities to validate and discover cancer-related somatic mutations. Solving the latter allows for more confident identification of different clonal lineages within a single tumor, which could inform understanding cancer evolution as well as targeted treatment decisions.

Bulk sequencing of pancreatic ductal adenocarcinoma (PDAC) is particularly problematic because of the high stromal content and low tumor cellularity which further lowers variant calling sensitivity^4–6^. Present solutions include multi-regional sampling to increase sensitivity for variants with low allele frequency^7,8^ or laser-capture tissue microdissection to enrich for tumor content^9,10^, but they are laborious and not amenable to high-throughput. With targeted single-cell sequencing, because each cell is partitioned and PCR amplified individually, high-quality genomic data from a low percentage of tumor cells could potentially be extracted from the background noise, making it valuable for genomic studies of PDAC.

To date, several high-throughput single cell partitioning systems have been developed, including microfluidic platforms, nanowells and microdroplets, which have resulted in several reliable single-cell DNA library preparation technologies^11^. Tapestri^12,13^, as a microdroplet-based, targeted sequencing approach, allows for high cell-throughput (up to 10,000 cells per sample) and high coverage depth (>80X) of genomic sites of interest, and is therefore suited for high-resolution studies of key genetic variants within diseases. However, its use so far has been limited to cell lines and liquid primary tumor samples, or a small batch of solid tumor tissues at a time^14–20^. While methods to quickly and effectively dissociate clean single nuclei suspensions from solid tumor tissues have been extensively tested for single-nucleus RNA-seq (snRNA-seq)^21^, they have not been applied for single-nucleus DNA-seq (snDNA-seq) usage.

We optimized a snap frozen tissue single nuclei extraction workflow that yielded high throughput in generating the resulting snDNA libraries. Importantly, the workflow takes < 30 minutes per sample with minimal manual labor, thus ideal for processing large batches of solid tumor samples. Coupling the snDNA data with bulk whole exome sequencing (WES) or whole genome sequencing (WGS) data generated on the same samples, we were able to uncover novel, single-cell clonal relationships among key driver mutations.

## Results

### Optimization of workflow to extract and store single nuclei from snap frozen tissue

We recognized the need for a nuclei extraction workflow that has reduced hands-on operation, sample resuspension times, and total processing time, all of which hinder scalability and could potentially cause between-sample inconsistency in quality (clumping, debris) and final yields. Thus, we used an automated nuclei extraction machine^22^ for homogenizing frozen tissues into single nuclei suspensions. The nuclei were then passed through a sucrose gradient to strip away debris before microdroplet encapsulation. The entire procedure takes ~30 minutes per sample and requires a single step of pelleting and resuspension (**Figure 1a; Methods)**. Although the resulting nuclei clumping % and nuclei concentration varied across samples of different starting sizes, cellularity and morphology, most primary pancreas tumor samples of volumes ⩾ 8 mm^3^, regardless of collection method (resection vs autopsy), resulted in final nuclei suspensions with ⩽ 10% clumping and >2,000 nuclei/ul suspended in > 35ul buffer as input for Tapestri (**Figure 1b)**. Exceptions were samples with extremely high fat/stroma content or random technical errors, which either limited input nuclei concentration or increased clumping % as observed by microscopy.

**Figure 1:**
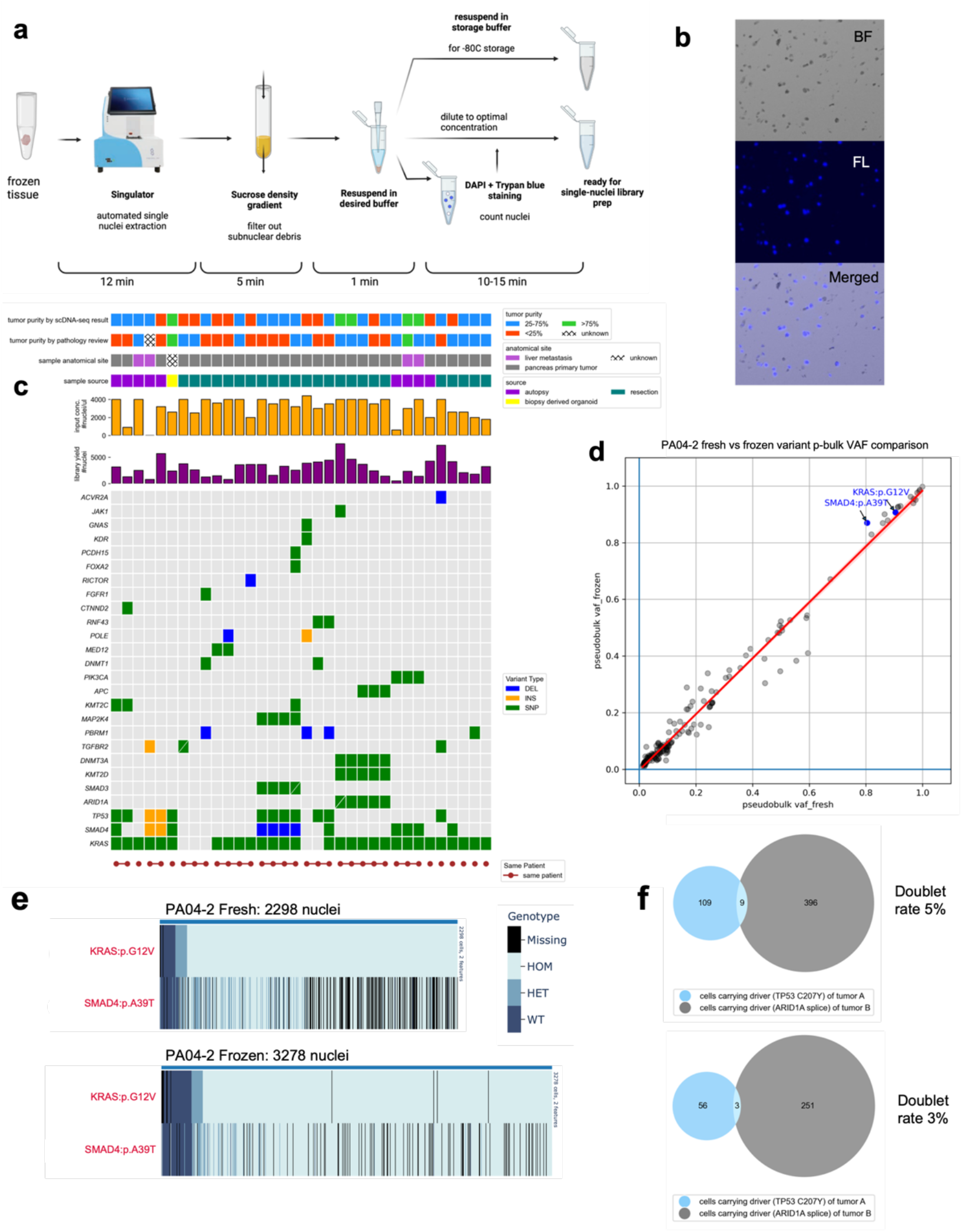
Frozen tissue single nuclei extraction workflow for snDNA-seq. **a.** Overview of frozen sample single nuclei extraction workflow for Tapestri snDNA-seq. **b.** Representative microscopic images of extracted single nuclei, stained with Trypan blue (brightfield, BF), DAPI (fluorescence, FL). **c.** Technical and genetic profile of each biologically distinct sample. A total of 38 samples were processed, 34 of which were biologically distinct samples from 16 unique patients. Genetic profiles were based on bulk sequencing. **d.** Pseudobulk (p-bulk) VAF comparison of all 160 shared variants between libraries prepared with fresh vs frozen (3 weeks) nuclei of sample PA04-2. Key drivers preidentified by bulk WES in this case are highlighted; regression line with 90% confidence interval is drawn. **e.** Single-cell genotype heatmap of snDNA-seq libraries generated by fresh vs frozen nuclei of sample PR04-2. Each row represents a bulk data-validated driver variant of this case, while each column represents a single nucleus in the library. The nuclei were sorted based on *KRAS* variant’s VAF in ascending order from left to right. **f.** Venn diagrams showing colocalization of genotypes belonging to two separate tumor cell populations in one snDNA-seq library (two replicates shown); the cells carrying both genotypes were identified as doublets. Nuclei suspension extracted from two tumor samples from different patients were mixed and subject to snDNA-seq. The two distinct tumor populations were identified by genotype for their respective driver variants *TP53 p.C207Y* and *ARID1A splice*.

With this protocol, we prepared 38 snDNA libraries from 34 biologically distinct snap frozen PDAC samples from 16 patients. These samples were purposely selected to represent primary tumors and metastases, different tissue collection methods, and with varying tumor purities (**Figure 1c; Supplementary Table 1**). Each was analyzed with a custom 186-amplicon panel covering 93 frequently mutated genes in PDAC (**Supplementary Table 2)**. The mean library yield of single nuclei with sufficient reads (**Methods)** was 2867 complete nuclei (standard deviation = 1672.67). For context, two previous studies^17,19^ using primary acute myeloid leukemia (AML) cell suspensions for Tapestri resulted in on average 5072 complete cells/sample (146 samples) and 6102 complete cells/sample (154 samples) in the final libraries.

We next determined the extent to which extracted single nuclei could be stored in suspension without affecting the yield. For two different samples (PA04-2 and PA04-3) we cryopreserved a portion of the extracted nuclei (**Methods**) then thawed these frozen nuclei suspensions after 3 weeks and 14 weeks respectively as input for snDNA-seq. This allowed us to compare both the nuclei morphology and the resulting snDNA-seq results between freshly extracted and cryopreserved nuclei of the same biological samples. We found that the freeze-thaw-resuspension process had >80% recovery rate and minimal change in nuclei morphology and clumping % (**Supplementary Figure 1a-b**). Moreover, the final library yield did not decrease when prepared with frozen nuclei; on the contrary, frozen nuclei gave slightly higher yield than fresh nuclei for both samples (**Supplementary Figure 1c**). Frozen nuclei generated similar quality results as freshly extracted nuclei as measured by sharing the majority of high-quality variants (**Methods**) (**Supplementary Figure 1d, e)**, and having largely linearly correlated pseudobulk VAF for all shared variants (**Figure 1d; Supplementary Figure 1f**). For both samples, fresh and frozen nuclei revealed highly concordant genotypes as well (**Figure 1e; Supplementary Figure 1g**). We also noted that for sample PA04-3 the frozen nuclei snDNA-seq result revealed a subclone not present in the fresh counterpart (**Supplemental Figure 1g**); this subclone was identified in both replicates of PA04-2, that was taken from a different region of the same liver metastasis. Such a difference could be attributed to different numbers of nuclei in the snDNA library (1371 vs. 635) between the frozen and fresh replicate of PA04-3, or simply technical variation.

While the multiplet rate inherent to the Tapestri scDNA library preparation method has been estimated to be 5-8% by its manufacturer^23^, we sought to estimate the multiplet rate associated with our entire workflow when using snap frozen tissue. We selected two samples from two distinct patients, each with a tumor population characterized by a distinct driver mutation (*TP53* p.C207Y vs. *ARID1A* splice) that was orthogonally validated by bulk WES as well as independent Tapestri runs. Similar sized pieces of each tissue sample were mixed together and subjected to the entire nuclei extraction workflow followed by scDNA library generation. Next, we assigned each cell to the two originating tumors based on its genotype for the two driver mutations. The number of barcodes that carried somatic variants from both tumors are as shown in the Venn Diagrams (**Figure 1f**). Using a mixture model (**Supplementary Note**) we derived the doublet rate to be 3-5%.

### Validation against bulk sequencing results

To validate the robustness of our method for detection of single-nucleotide variants (SNVs) (**Supplemental Figure 2**), we selected 18 samples previously used for bulk WES sequencing (**Methods**). When limiting to variants present within the targeted panel, a strong concordance was noted between unfiltered coding variants called from frozen nuclei versus those called by WES (**Figure 2a; Supplementary Figure 3a)**. When we next limited to filtered, high quality variants (**Methods**) there was virtual complete concordance (**Figure 2b)**. Moreover, despite technical differences inherent to WES versus Tapestri (input tissue slice, DNA library prep technology and sequencing depths) the bulk WES VAFs and snDNA-seq pseudobulk VAFs for all shared variants, except for those with ⩽ 2 alternative reads in bulk WES results, were linearly correlated for the majority of samples (**Supplementary Figure 3b-d**).

**Figure 2:**
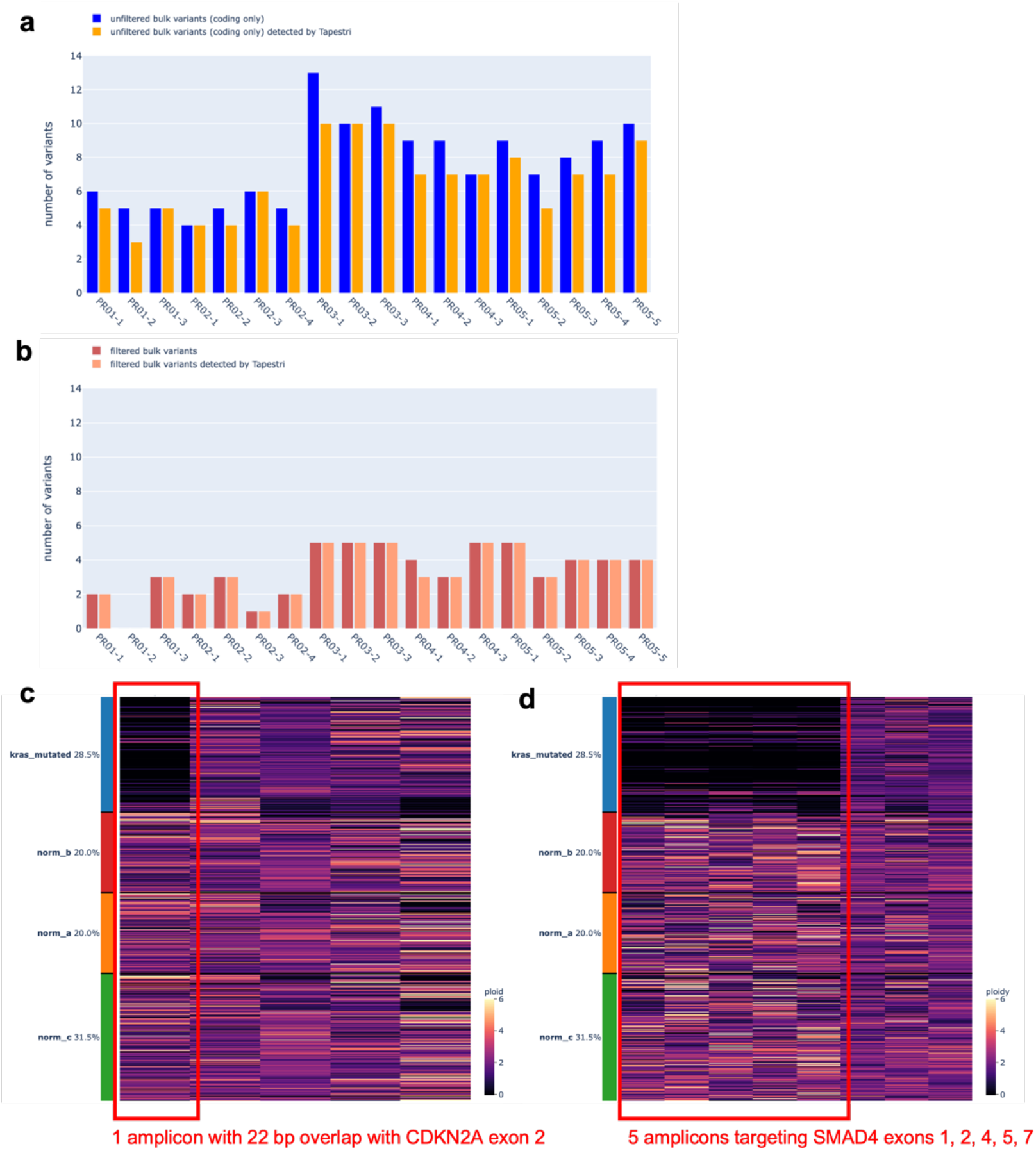
snDNA-seq was able to capture SNV and CNV pre-identified by bulk-sequencing with high sensitivity. **a.** For 18 samples of the same bulk WES sequencing cohort, histogram showing the number of unfiltered (Methods), coding variants called by bulk (blue) and among them, the number detected by snDNA-seq (orange). **b.** For the same samples, histogram showing the number of filtered coding variants called by bulk (dark red) and among them, the number detected by snDNA-seq (light red). **c-d.** Single-cell per-amplicon ploidy heatmap of select amplicons covering chromosomes 9 (c) and 18 (d) of sample PA02-1. Each row represents one cell while each column is one amplicon. Cells are divided into *KRAS* mutated group (blue) and 3 normal groups (red, yellow, green) (**Methods**) and hierarchically clustered within group. The amplicons spanning *CDKN2A* and *SMAD4* genes are outlined in red.

We next sought to determine if our scDNA-seq approach can detect subclonal copy number variations (CNVs) that may be difficult to call with low-depth/low-coverage bulk sequencing data. For this analysis we used sample PA02-1 with known homozygous deletions of *SMAD4* and *CDKN2A*^24^. By calculating the single-cell per-amplicon (~200bp) ploidy based on read counts (**Methods**), we were able to identify both homozygous deletions (ploidy~0) in the neoplastic population despite it comprising 28.5% of all cells (**Figure 2c, d; Supplementary Figure 4a-d)**.

### scDNA-seq is applicable to low-cellularity, low-tumor content samples

Low cellularity and tumor content in certain clinical settings, such as with fine needle aspirations, core needle biopsies or for PDAC in general^25^, represent major hurdles for bulk-sequencing and hinder the quality of resulting genomic information. We therefore tested our workflow’s performance in such settings.

We identified one sample PA04-1 of a primary pancreas tumor with both low tumor cellularity (neoplastic cells occupying <50% area of tissue section) and low overall cellularity due to a high fat content (**Figure 3a**). Despite use of a tissue sample of similar size (~8mm^3^) as others studied, we extracted a total of 30,000 nuclei, much lower than the optimal number of 200,000 as input for Tapestri. Ultimately, we captured only 479 nuclei in the final library, approximately 6-fold lower than average. Nonetheless, both driver gene variants (*KRAS p.G12V*, *SMAD4 p.A39T*) identified by bulk WES of this same sample were identified with high quality read data (**Supplementary Figure 5a, b**) and indicated 113 likely tumor cells (23.6% of all) captured for this sample based on the presence of a clonal *KRAS* variant (**Figure 3b, c)**.

**Figure 3:**
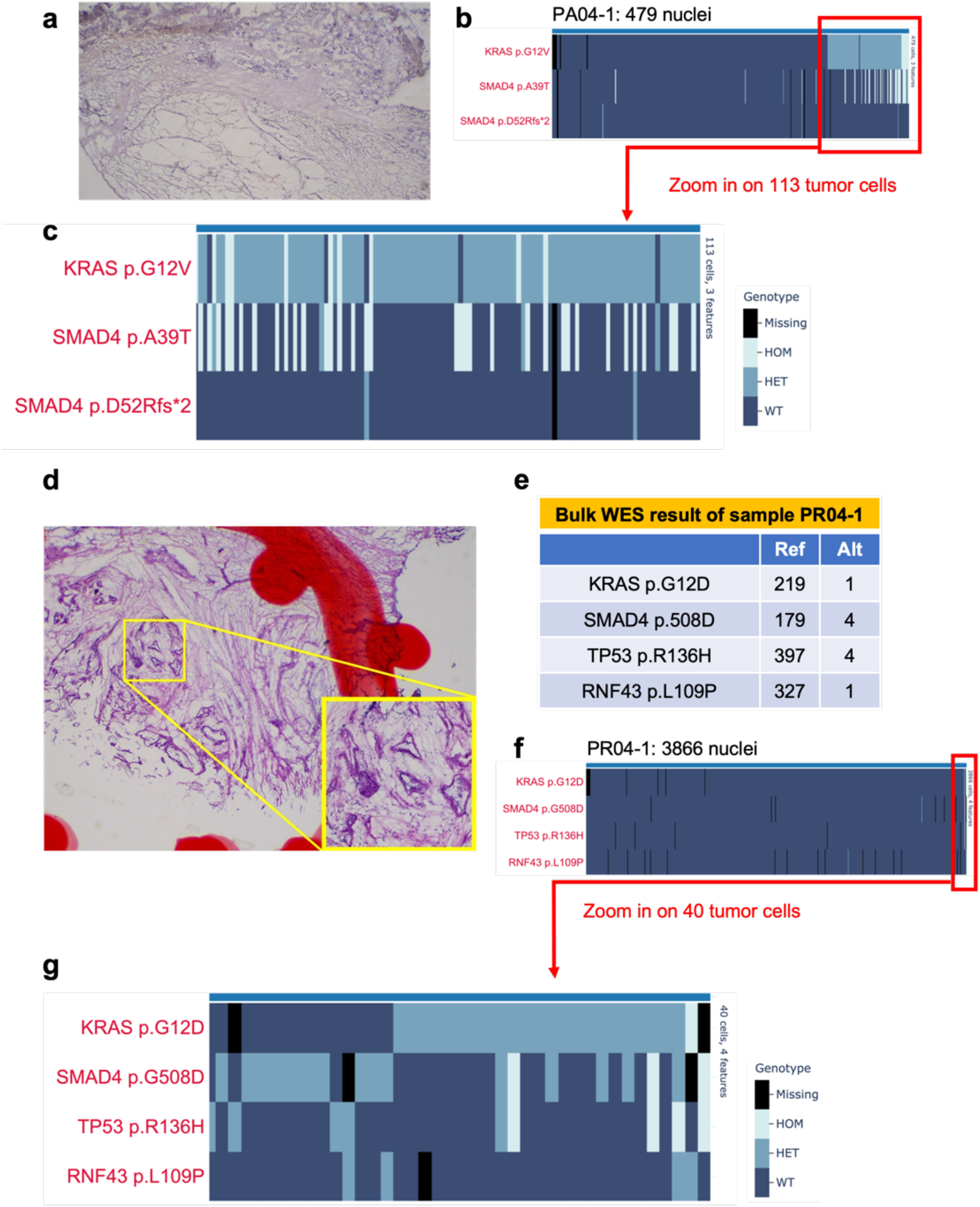
Tapestri snDNA-seq’s performance in limiting settings. **a.** Representative Hematoxylin and eosin (H&E) stained histology image of sample PA04-1, a sample of extremely low cellularity due to high fat content. **b.** Single-cell genotype heatmap of sample PA04-1. A total of 479 captured single nuclei were sorted based on *KRAS* variant allele frequency (VAF) in ascending order from left to right. **c.** Single-cell genotype heatmap of sample PA04-1, zoomed in on 113 putative tumor cells, defined by the presence of *KRAS p.G12V/SMAD4* p.39T mutation. **d.** Representative H&E histology image of sample PR04-1. **e.** Bulk WES result of sample PR04-1. **f.** Single-cell genotype heatmap of PR04-1. A total of 3866 captured single nuclei are sorted based on *KRAS* VAF in ascending order from left to right. Tumor cells, defined by the presence of *KRAS* p.G12D/*SMAD4* p.G508D/*TP53* p.R136H mutations, cluster to the right and are barely visible. **g.** Single-cell genotype heatmap of PR04-1, zoomed in on 40 putative tumor cells.

With a second sample of extremely low tumor purity as revealed by pathology review (**Figure 3d**) and bulk sequencing (**Figure 3e**), Tapestri data identified 40 out of 3866 nuclei (1%) of this sample carrying at least one of *KRAS p.G12V*, *TP53 p.R186H*, *or SMAD4 p.508D* (**Figure 3f, g)**. Again, the driver variants were genotyped with high quality read data (**Supplementary Figure 5c, d**). Intriguingly, the single-nucleus colocalization of the three main drivers violated the assumption of the infinite sites model: more than half of the nuclei carry the *KRAS* mutation and among them, a subset carry *SMAD4* and *TP53*, yet a significant number of nuclei were wildtype for *KRAS* yet mutated for *SMAD4/TP53*. If it were assumed that *KRAS* were mutated first and *SMAD4* and *TP53* mutations followed, a likely explanation would be that a subset of tumor cells lost their mutant *KRAS* allele through loss of heterozygosity (LOH). Although allelic dropout (ADO) inherent to the library preparation method might also factor in, similar patterns observed in snDNA-seq results of two other samples of this case (M12-2, M12-3) seemed to support the abovementioned theory (**Supplementary Figure 5e, f**). However, more rigorous statistical modeling is required for validation.

### snDNA-seq identified two mutually exclusive clones bearing two different KRAS mutations in the same PDAC patient

For a PDAC surgical resection case PR02, based on both MSK-IMPACT sequencing (high-depth targeted sequencing) and bulk WES, we identified the major tumor clone carry the hotspot *KRAS* p.G12D mutation; hints of a minor *KRAS* p.G12V clone existed but were on the borderline of the technologies’ detection sensitivity (**Figure 4a**). Single-nucleus genotype heatmaps and Venn diagrams (**Figure 4b-e**) of multiregional samples PR02-3 and PR02-4 suggested colocalization of the major *KRAS p.G12D* with another likely driver *TP53* p.C203Y, which signified the major tumor clone in this sample; the minor *KRAS* p.G12V-bearing clone was mutually exclusive with the above two drivers and did not colocalize with any known driver gene mutation at similar clonal frequency. In PR02-4, while the major clone consisted of 221 cells (6.12% of all cells), the minor clone was only 12 cells (0.33% of all cells, **Figure 4e**), further buttressing the technology’s sensitivity. The minor *KRAS* p.G12V clone in both samples was substantiated with high quality read data (**Supplementary Figure 6a-b**); digital droplet PCR (ddPCR) also proved the presence of the *KRAS* p.G12V (**Supplementary Figure 6c**). Pathology review did not identify any apparent secondary neoplastic or metaplastic structure (**fig 4f)**. This observation aligns with several other studies^6,26^ in suggesting that multiple *KRAS* genetic variants may coexist in one patient’s PDAC precursor/tumor.

**Figure 4:**
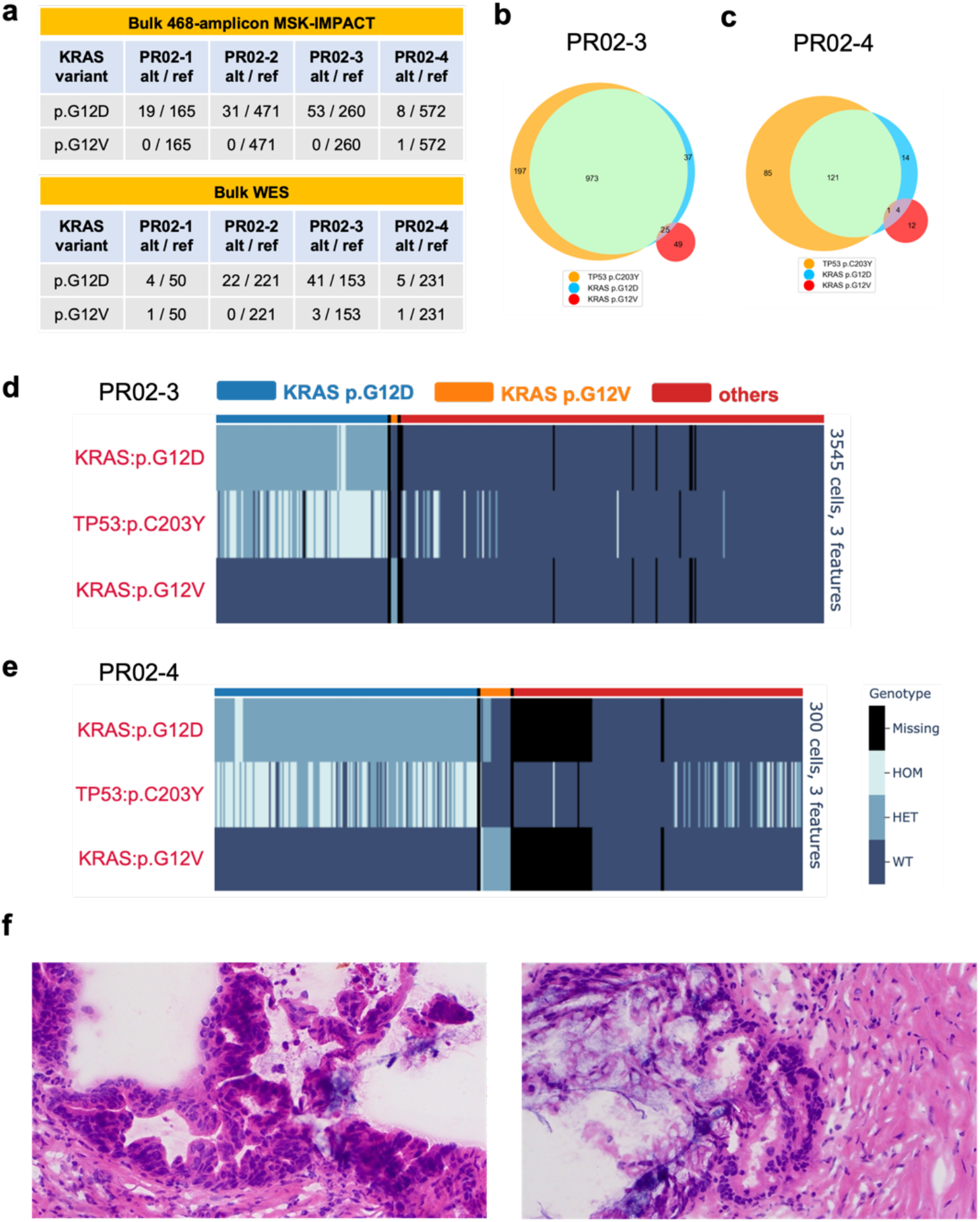
snDNA-seq identified two mutually exclusive clones bearing two different KRAS mutations in one pancreatic cancer patient. **a.** Bulk-sequencing calls of *KRAS* variants of patient PR02’s 4 multiregional primary tumor samples. **b-c.** Venn diagram showing colocalization pattern of genetic variants *TP53* p.C203Y, *KRAS* p.G12D, *KRAS* p.G12V in single nucleus of samples PR02-3 (**b**), PR02-4 (**c). d-e.** single-cell genotype heatmap of samples PR02-3 (**d**), PR02-4 (**e,** zoomed in on 300 cells where tumor cells cluster). The *KRAS* p.G12D & p.G12V clone identities (above heatmaps) are identified as cells having “HET” or “HOM” genotype of each variant and are colored as labeled. Cells are hierarchically clustered based on the two KRAS variants’ single-cell AF. *KRAS* p.G12V clone size was 57 cells in PR02-3, 17 cells in PR02-4. **f.** representative H&E histology images of sample PR02-3.

### snDNA-seq identified complex clonal structures in a KRAS-WT PDAC

The *KRAS* gene is mutated in >90% of all PDAC’s and signifies the phenotype of MAPK-ERK pathway hyperactivation^25^. By bulk exome sequencing of three spatially distinct samples of resected PDAC PR01, we noted that this case was wild type for *KRAS* yet contained an *FGFR1* p.T50K mutation as well as two distinct *TGFBR2* mutations (p.M450I, p.A451G) on the same allele. Pathology review of the samples’ H&E slides identified two well-isolated populations of PDAC cells with distinct histological features. A population of PDAC cells characterized as dilated glands with extensive stroma was exclusively present in sample PR01-1 (**Figure 5a, left)**, while another population characterized as small nests of tumor cells was exclusively present in PR01-2 **(Figure 5a, right)**. Sample PR01-3 had both populations present in the same tissue section. We performed snDNA-seq of all 3 samples and discovered that the *FGFR1* and *TGFBR2* mutations corresponded to two mutually exclusive clones: sample PR01-1 had only the *TGFBR2* double-mutated clone (**Supplementary Figure 7a**), PR01-2 only the *FGFR1* mutated (**Supplementary Figure 7b**), while PR01-3 contained both clones that were mutually exclusive at the single-cell level (**Figure 5b-c, Supplementary Figure 7c**). To determine the extent to which these two clones were unique neoplasms versus subclones that shared a common ancestor, we included all other high-quality coding & non-germline variants to define a putative normal cell population and computed the median per-amplicon ploidy within each clone (**Methods**). This revealed many shared CNVs between the two clones that included allelic losses of *ARID1A, TGFBR, FGFR1* and *SMAD4* as well as a homozygous deletion of *CDKN2A* **(Figure 5d, Supplementary Figure 7c)**. Collectively these data suggest that large-scale copy number aberrations preceded the formation of the two distinct SNV clones.

**Figure 5:**
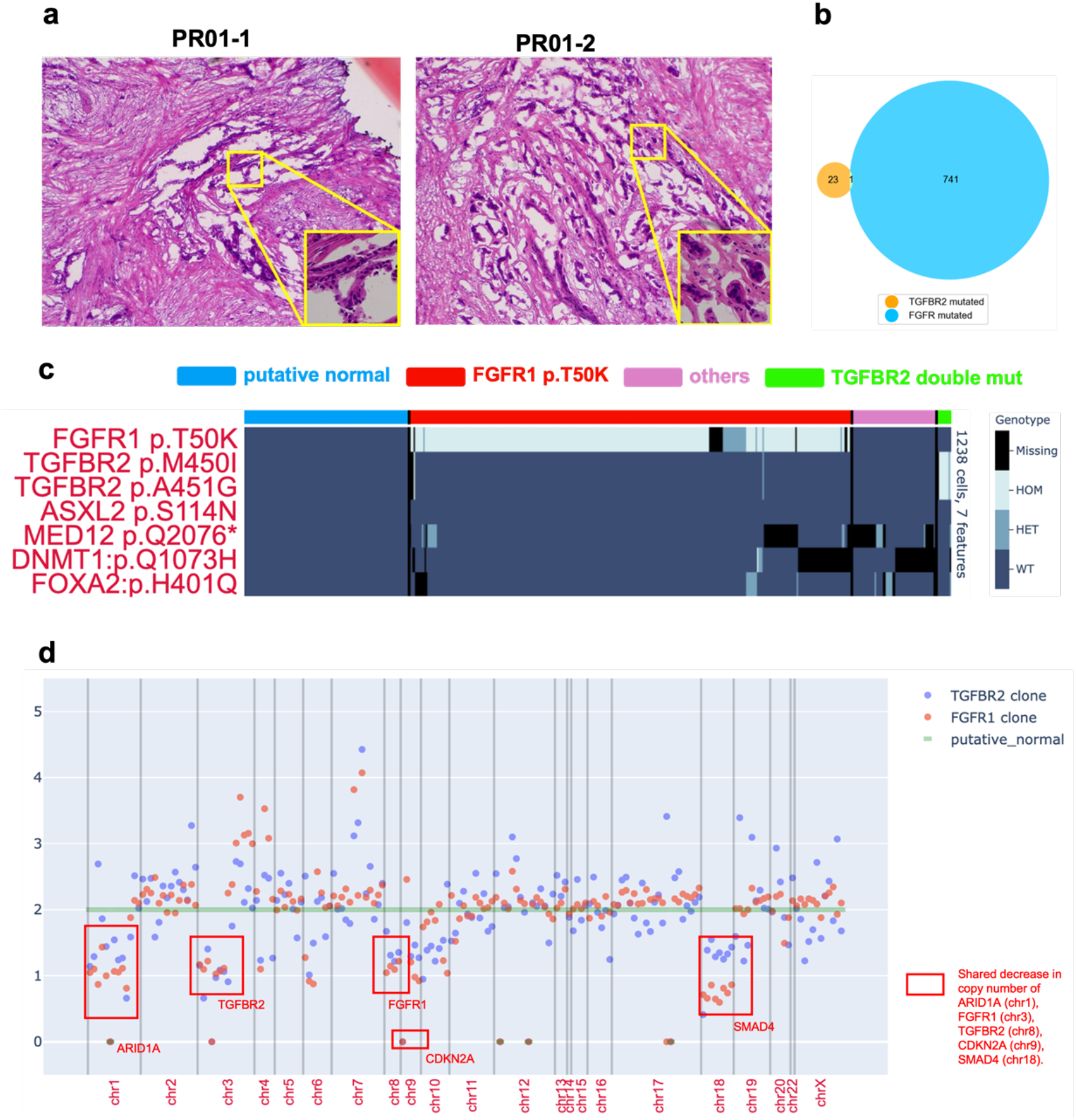
snDNA-seq identified two mutually exclusive SNV clones in a PDAC patient without *KRAS* mutation. **a.** Representative H&E histology images of PDAC regions in samples PR01-1 (**left**, only the *TGFBR2* double mutation was present), PR01-2 (**right,** only the *FGFR1* mutation was present). **b.** For sample PR01-3 (both the *TGFBR2* double mutation and *FGFR1* mutation were present), Venn diagram showing single-cell colocalization pattern of the *TGFBR2* double mutation and *FGFR1* mutation. **c.** Single-cell genotype heatmap of 7 important variants pre-identified by bulk WES for sample PR01-3. Each cell’s clone identity (color strip above heatmap) is colored as shown by the figure legend. The *FGFR1* and *TGFBR2* SNV clones are defined as cells with non-WT genotype of each gene. The “putative normal” clone is identified as cells with “WT” genotype of all 7 genetic variants. Cells are hierarchically clustered within each clone. **d.** Median per-amplicon ploidy of the *TGFBR2* double mutation clone, the *FGFR1* mutation clone and the putative normal clone found PR01-3. The putative normal clone is set as diploid baseline (green); amplicons with notable copy number loss and their corresponding genes are labeled.

### snDNA-seq revealed step-wise evolution during PDAC metastasis

Through snDNA-seq of three samples of PA04 (one primary tumor sample, two liver metastases) we identified sequential steps leading to TGF-β inactivation in association with cancer progression. Specifically, we identified five distinct subclones based on the genotypes of two genes, *KRAS* and *SMAD4* (**Figure 6a,** figure legend). These subclones were present in different proportions in each sample. Clone 1 (indicated by red) was characterized by a heterozygous *KRAS* p.G12V mutation, whereas clone 2 (indicated by orange) contained a *SMAD4* p.A39T mutation in addition to the *KRAS* mutation (**Figure 6c,d, Supplementary Figure 8a,b**). Clone 3 (light green) contained a greater degree of allelic imbalance for mutant *KRAS* in part due to loss of the wild type *KRAS* allele, best appreciated in the liver metastases (**Figure 6b-e; Supplementary Figure 8b,c**). Moreover, in PA04-3 specifically (**Figure 6d**), clone 3 contains two populations: one heterozygous for *SMAD4* p.A39T and one homozygous for this variant in association with allelic loss of part of the gene, suggesting that the missense mutation preceded the loss of heterozygosity event. Clone 4 (dark green) is characterized by a second mutation in *SMAD4* at p.D52Rfs*2, and clone 5 (dark blue) illustrates complete loss of both *SMAD4* mutations and hence complete homozygous deletion of the gene (**Figure 6b,c,e; Supplementary Figure 8b**).

**Figure 6:**
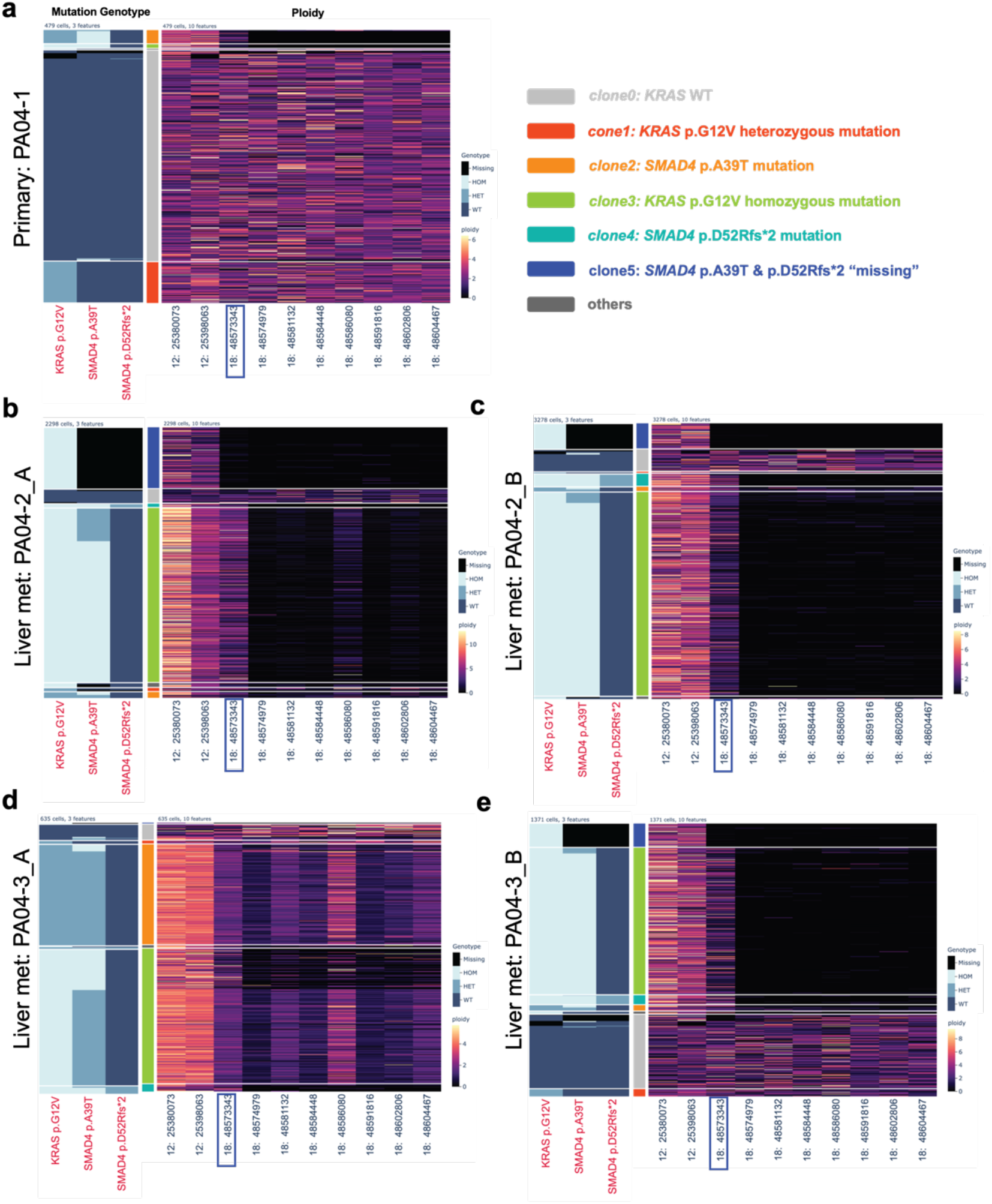
snDNA-seq revealed step-wise evolution during PDAC metastasis. **a-e.** Combined genotype and ploidy analysis of 3 multiregional samples of PDAC autopsy case PA04. In each panel, single-cell genotype heatmap (of important *KRAS* and *SMAD4* variants preidentified by bulk WGS) is placed on the left, each cell’s clone identity (as defined by *KRAS* and *SMAD4* genotype as shown in the legend) in the middle and single-cell per-amplicon ploidy heatmap on the right. Cells (rows) are sorted hierarchically within each clone. Each panel corresponds to PA04-1 (**a,** primary tumor), PA04-2_A **(b,** liver met slice 1, fresh nuclei), PA04-2_B **(c,** liver met slice 1, frozen nuclei), PA04-3_A (**d,** liver met slice 2, fresh nuclei) and PA04-3_B (**e,** liver met slice 2, frozen nuclei). For the ploidy heatmap, each amplicon’s starting genomic location is labeled on the x-axis. The amplicon where the *SMAD4* p.A39T and p.D52Rfs*2 mutations took place is outlined in blue.

Based on these observations, we conclude that four of five clones were pre-existent in the primary site and disseminated to the secondary site, either directly from the primary tumor or by way of another unsampled secondary site. It is ambiguous whether the fifth clone with homozygous deletion of most *SMAD4* exon arose in the primary site or not, since it was not present in the one primary sample we sequenced. While differences in clone proportions may indicate selection for specific genotypes (i.e. KRAS amplification, SMAD4 deletion) we cannot rule out stochastic events during sample preparation that may have enriched for some nuclei over others (**Figure 7**). Perhaps most notable is the finding of five distinct events affecting *SMAD4:* p.A39T mutation, LOH, partial homozygous deletion, D52Rfs*2 mutation and finally homozygous deletion of the remaining exonic region containing these two mutations. This finding is in keeping with inactivation of cell-intrinsic TGFβ signaling as a critical aspect of PDAC metastasis^27–30^, as well as ongoing clonal selection for survival benefits^25,31^.

**Figure 7:**
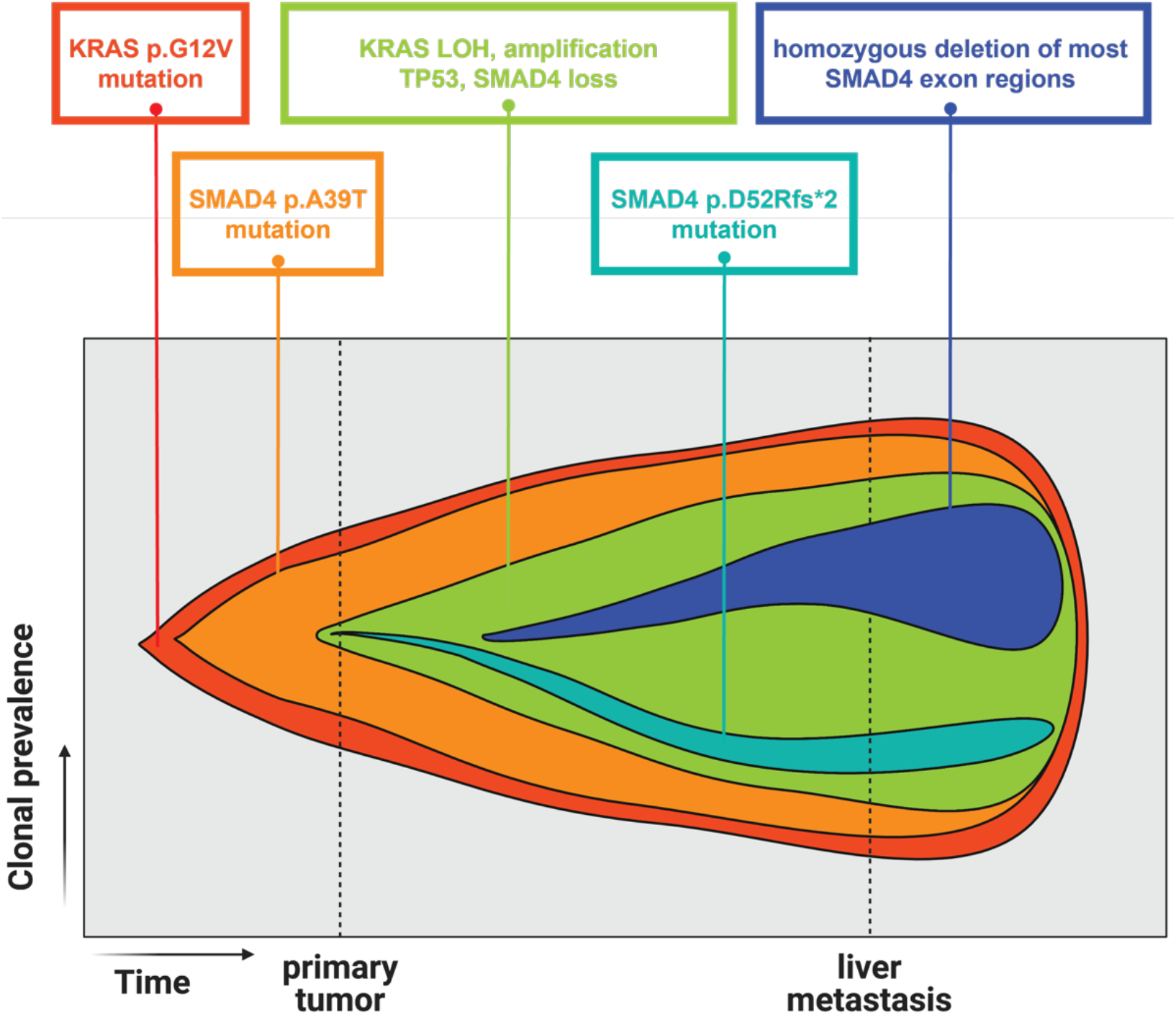
Stepwise evolution pattern of PDAC case PA04. Fishplot demonstrating the stepwise evolution of PDAC case PA04, as inferred by snDNA-seq results of three multiregional samples from the same patient. Relative evolution time was plotted on the x-axis and clonal prevalence on the y-axis. The two timepoints (primary, liver met) are arbitrarily labeled based on sample locations. In the primary tumor, cells first gained *KRAS* p.G12V mutation and then *SMAD4* p.A39T mutation; followed by *KRAS* allelic imbalance and *SMAD4* allelic loss. A second *SMAD4* mutation (p.D52Rfs*2) occurred in tandem to the first one in the primary tumor, followed by homozygous deletion of the entire *SMAD4* gene; this latter clone (blue) achieved high clonal fraction in the metastatic samples.

## Discussion

In this study, we used two commercially available products to assemble a highly automated workflow to generate Tapestri snDNA-seq libraries from snap frozen patient tissues. The workflow is fast, efficient and can be applicable for high-throughput clinical and translational research. Additionally, we recognized the value of storing excess single-nuclei suspensions for later use and verified a corresponding workflow that was free from issues such as nuclei quantity loss, nuclear envelope damage, or nuclei clumping. This workflow further illustrates the ability to maintain information pertaining to relative VAFs compared to matched WES data. Furthermore, while our custom panel was not designed for identification of CNVs, specifically homozygous deletions, we demonstrate the proof of principle that these events could be identified with confidence. These encouraging results, combined with the added information gleaned pertaining to co-occurring and mutually exclusive genetic events, suggests this technology and workflow is ideally suited to settings in which samples sizes are small or limited.

A caveat of this snDNA-seq technology is that albeit its high cell-throughput, it is a targeted approach that focuses on a pre-designed panel of genes which might be insufficient for certain research questions that require unbiased study of the cancer exome/genome^32–35^. This also implies a general challenge of single-cell research: because the total library size is limited by sequencing cost and data processing capacities, there is always a trade-off between the total cell-throughput and the amount of information one can extract from each single cell. Another caveat, in this instance related to our computational analysis, is that the current variant calling pipeline for Tapestri applied GATK HaplotypeCaller and enabled default read downsampling which might be suboptimal for such high-depth single-cell data in a cancer setting. We did not apply matched normal/panel of normal (PON)-aided filtering in our pipeline; a benefit of this approach is that it enabled us to see many germline variants that can be used for quality control or phylogeny modeling, although this benefit is balanced by outputs containing a large number of artifacts. Ultimately, to enable novel somatic variant discovery a more robust variant calling pipeline is needed to adapt to the noise profiles particular to this PCR-based high-throughput single-cell DNA sequencing technology.

The field of single-cell transcriptomics began earlier than single cell genomics, and accompanying analysis methods have been flourishing in the past 5 years^36,37^. Single-cell transcriptome-oriented methods, such as clustering or gene-set enrichment analysis, are generally not optimal for single-cell genomic data, because the cell-cell difference for the latter is minimal in comparison, often only on a handful of genetic mutations. The best method would be to rely on the ancestor-descendent relationship between every pair of cells’ genomic sequence and build single-cell phylogenies. Several models for the evolution of SNVs in cancer have been developed to date^38–44^, but only two^43,44^ have been constructed to account for the frequent, complex aneuploidy in cancer, one of which is suited for the high throughput and depth of the dataset presented in this paper^44^. From the combined SNV and CNV data of case PR01’s multiregional samples (**Supplementary Figure 7a-c)**, we could already see many complex clonal structures than could not be defined by driver SNV clones alone; there was apparent large-scale aneuploidy that was ancestral to the driver SNVs as well as focal loss of genomic segments in descent cells. From analyzing multiregional autopsy samples of case PA04, we also surmise that manual investigation of subclonal structure based on driver genes could easily fall short in uncovering more granular evolution patterns. Efficient methods to analyze such datasets, potentially leveraging matched bulk sequencing data for greater coverage of the genome, need to be developed and tested in order to fully unleash the power of single-cell genomic studies on multiregional samples.

## Supporting information

Supplementary_Table 2- Tapestri panel coverage of canonical exons

Supplementary_Table 1- sample profiles

Supplementary_Document-Doublet_estimation

## Data availability

Sequence data have been deposited at the European Genomephenome Archive (EGA), which is hosted by the European Bioinformatics Institute and the Centre for Genomic Regulation, under accession number EGAS00001006024. Further information about EGA can be found at https://ega-archive.org and “The European Genomephenome Archive of human data consented for biomedical research” (http://www.nature.com/ng/journal/v47/n7/full/ng.3312.html). All data supporting the findings of this study are in the process of uploading to EGA to be available upon publication.

## Methods

### Ethics statement

Use of samples used in this study was approved by IRB review at Memorial Sloan Kettering Cancer Center.

### Patient sample collection and preprocessing

Patient samples used in this study mainly consist of two categories-multiregionally sampled surgical resection of primary pancreatic cancer, and multiregionally sampled autopsy of metastatic pancreatic cancer. Detailed patient, sample information is summarized in **Supplementary Table 1**.

For multiregional sampled surgical resections: treatment naïve patients with tumors ≥2cm on cross-sectional imaging were identified preoperatively. A single cross-sectional piece of tumor was sampled sequentially using a cartesian coordinate system with 0.6cm x 0.6cm grid, with 3 to 5 samples obtained from each tumor. Adjacent normal pancreas or duodenum was also collected. All samples were stored at −80 degrees Celsius until use.

For multiregional sampled autopsy: Tissues from three patients were used. All patients had a premortem diagnosis of PDAC based on pathological review of resected biopsy material and/or radiographic and biomarker studies.

Tissue sections were cut from tissue blocks embedded in optimal cutting temperature (OCT) compound, stained with hematoxylin and eosin (H&E) and reviewed by a gastrointestinal pathologist (S.U.) to estimate total cellularity, tumor purity and tissue quality. Normal samples were reviewed to confirm that no contaminating cancer cells were present.

### Bulk WES, WGS library preparation, sequencing and variant calling

Genomic DNA was extracted from each tissue using the phenol-chloroform extraction protocol or QIAamp DNA Mini Kits (Qiagen). WGS, WES and alignment were performed by the Integrated Genomics Operation and the Bioinformatics Core at Memorial Sloan Kettering Cancer (New York, NY). Briefly, an Illumina HiSeq 2000, HiSeq 2500, HiSeq 4000 or NovaSeq 6000 platform was used to target sequencing coverages of >60× for WGS samples and >150× for WES samples.

Sequencing reads were analyzed in silico to assess quality, coverage, as well as alignment to the human reference genome (hg19) using BWA. After read de-duplication, base quality recalibration and multiple sequence realignment were completed with the PICARD Suite and GATK v.3.1; somatic single-nucleotide variants and insertion–deletion mutations were detected using Mutect v.1.1.6 and HaplotypeCaller v.2.4. Such a process generates the “filtered” variant list for every sample. Then, all variants of all samples of the same sequencing cohort were pooled as a single list. Each sample’s BAM file were used to compute “fillout” values (total depth, reference allele counts, alternative allele counts) for each variant in the pooled list. This process aimed to rescue variants that were detected with high confidence in multiregional sample #1 but with low confidence in multiregional sample #2 of the same patient; the output corresponded to the “unfiltered” variant list.

### Nuclei extraction from frozen tissue, counting, QC, sorting and cryopreservation

Single nuclei from OCT-embedded snap frozen primary tissue samples were extracted using the Singulator 100 machine (S2 Genomics) with its extended nuclei dissociation protocol. After extraction, nuclei solution was centrifuged at 800g for 5 minutes in a swing bucket with a reduced braking in a 0.25M Nuclei PURE Sucrose solution (Sigma-Aldrich) to filter out debris.

Nuclei were stained with Trypan blue and manually inspected under a brightfield microscope for clumping%, which was estimated as the number of clumped particles out of all single particles within one field of view. Nuclei concentration was estimated by DAPI staining on a Countess II FL automated cell counter. Clumping% and nuclei concentration were both measured ⩾ 2 times for each sample. A final concentration of 4000 nuclei/ul suspended in 50ul Mission Bio cell buffer was targeted per sample prepared.

After up to 200,000 nuclei were taken for Tapestri library preparation, the remaining nuclei were resuspended in Sigma Aldrich Nuclei PURE storage buffer and immediately frozen on dry ice, before being transferred to −80C freezer for long-term storage. If needed, nuclei were thawed on ice until the solution was clear, and centrifuged with the same settings as described above for pelleting and buffer exchange.

### Single-nuclei library preparation and sequencing

Nuclei were suspended in Mission Bio cell buffer at a maximum concentration of 4000 nuclei/ul, encapsulated in Tapestri microfluidics cartridge lysed and barcoded. Barcoded samples were then put through targeted PCR amplification with a custom 186-amplicon panel covering important PDAC mutational hotspots in our sample cohort (**Supplementary Table 2**). PCR products were removed from individual droplets, purified with Ampure XP beads and used as templates for PCR to incorporate Illumina i5/i7 indices. PCR products were purified again, quantified with an Agilent Bioanlyzer for quality control, and sequenced on an Illumina NovaSeq.

### Single-nuclei DNA library quality control, cell-calling and variant calling

FASTQ files for single-nuclei DNA libraries were processed through Mission Bio’s Tapestri pipeline with default parameters. Briefly, it trims adaptor sequences, aligns reads to the hg19 genome (UCSC), assigns reads to cell barcodes. The CellFinder module then filtered for barcodes corresponding to “complete cells/nucleus” based on total read completeness (>8 * number of amplicons) and per-amplicon read completeness (>80% data completeness for working amplicons, which are defined as amplicons with > 0.2*mean of all amplicon reads per qualified barcode). It next used GATK HaplotypeCaller to call variants individually on each cell, and then GATK GenotypeGVCFs to jointly genotype all cells using genotype likelihoods from the previous step. The unfiltered VCF was parsed into an HDF5 file containing single-cell variant and per-amplicon read count matrices compatible with downstream analysis.

### Single-cell genotyping and cell-variant pair filtering

The HDF5 file output from above was analyzed mainly by Mission Bio’s python-based analysis package Mosaic, with a modified genotyping and variant filtering module. As shown in **Supplementary Figure 1**, with a single-cell variant call matrix, we started by assigning a genotype to each cell-variant pair. First, we defined the minimum depth at 5 reads and any variant in any cell with depth below the threshold in a cell would be assigned as “missing”.

Then, we used cutoffs:

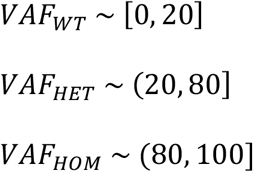

to assign each variant’s genotype (WT-wildtype; HET-heterozygously mutated; HOM-homozygously mutated) in each cell, thus allowing 20% of reads of a barcode to be false positives potentially caused by barcode contamination.

Finally, we set the threshold for the alternative (mutant) read count to 3 reads to convert low-quality heterozygous calls back to wildtype to arrive at the final cell-variant genotype matrix.

For the sake of comparing the same sample under different conditions, we computed a list of high-quality variants for each library as follows: we first discarded variants that have “missing” genotype in more than 75% cells (while whitelisting certain genes that are known to be prone to homozygous deletion in PDAC, such as *SMAD4*, *CDKN2A*). Through inspecting the distributions of cellular prevalence of variants across different read depths and total cell numbers, we determined that a mutational prevalence of 0.5% is a feasible and effective cutoff to filter out most technical artifacts. Any variant mutated in more than 0.5% of all cells was added to the high-quality variant list. Worth mentioning is we still observed obvious artifacts in these final lists, which need to be addressed with more rigorous filtering in further studies.

### Doublet model and calculation

Please see attached **Supplementary Document**.

### Single-cell per-amplicon ploidy calculation

The ploidy calculation was mainly based on Mission Bio’s Mosaic package. The per-amplicon read counts were normalized first within the same cell across different amplicons by mean read depth, and then within the same amplicon across different cells by median read depth. Note the median read depth across different cells only considered good-quality cells, which are defined as those with at least 1/10 number of reads as that of the cell with the 10^th^ rank in terms of read count.

Then the per-amplicon ploidy was calculated by setting a group of cells as diploid baseline based on *a priori* knowledge (e.g. *KRAS* mutational status) and taking the ratio of every other cell’s per-amplicon read count against that group’s per-amplicon median read count.

To test the robustness of our ploidy calculation, we picked one sample with known *KRAS* mutation and cancer-related aneuploidy based on bulk sequencing and validated with DNA microarray. We started by separating a scDNA-seq library into two groups-*KRAS*-mutated and *KRAS*-WT, with the latter assumed as mostly normal cells. Then we divided the normal cell population randomly into 3 groups (norm_a, norm_b, norm_c), used one group (norm_a) as diploid baseline and calculated other groups’ ploidy against it. As shown in the **Supplementary Figure 3**, the two other putative normal cell groups had their median per-amplicon ploidy aligning close to 2, which validates the diploid-defining rule; the *KRAS*-mutated group had apparent aneuploidy across most amplicons and *CDKN2A* and *SMAD4* loss, which validated our ploidy calculation.

For case PR01, because there was not an *a priori* clonal oncogenic driver such as a *KRAS* variant to be reliably used to determine a diploid population, we used 7 (including 2 *TGFBR2* mutation 2 bp apart) variants pre-identified by bulk WES to set up a rule: a cell with “WT” genotype for all 7 variants can be called putative normal and be used as diploid baseline.

---

**Supplementary figure 1:**
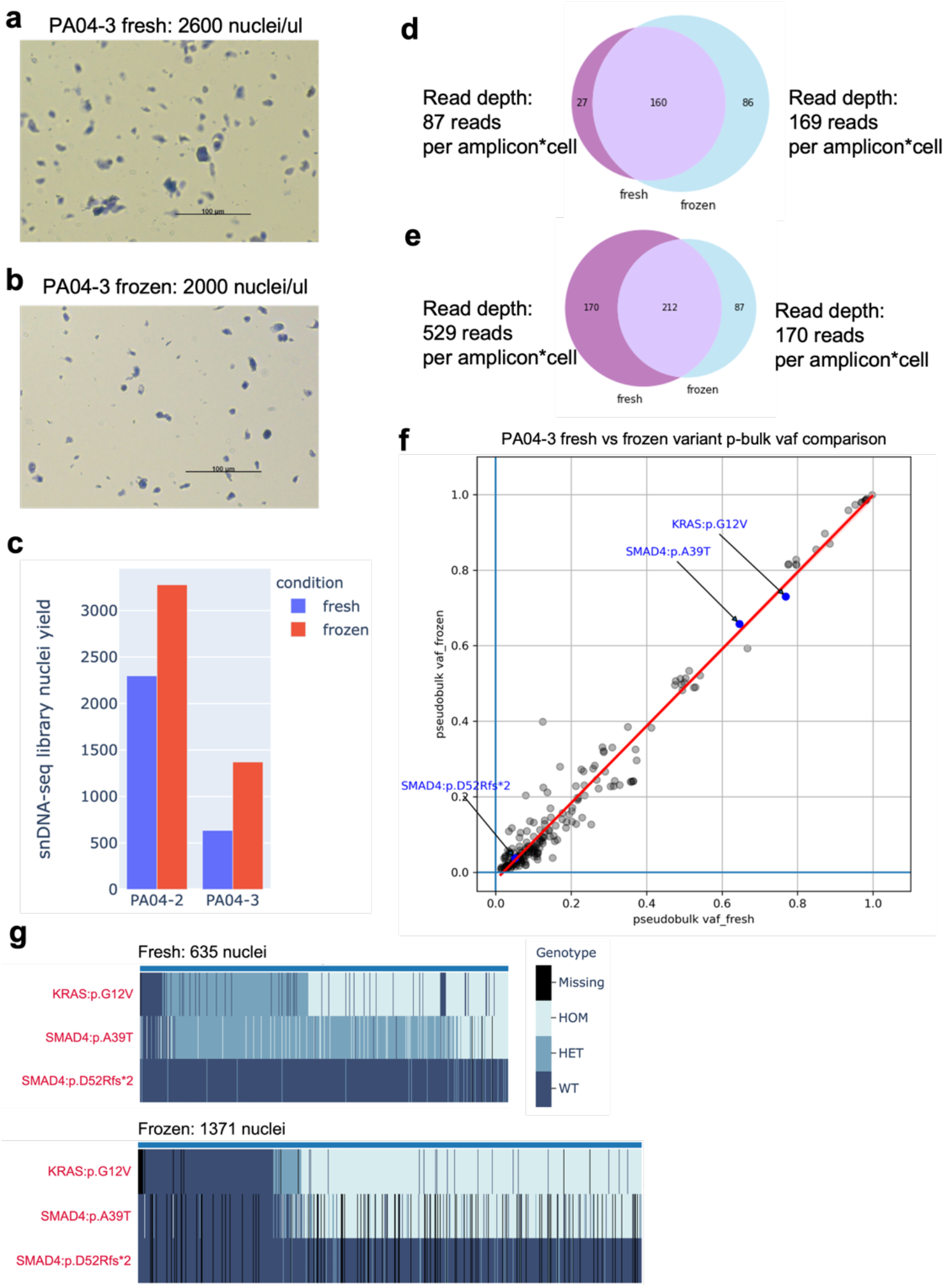
Nuclei cryopreservation results. **a-b.** Sample PA04-3 nuclei suspension before **(a)** and after cryopreservation **(b).** Nuclei were stained with Trypan blue and visualized under brightfield microscope. **c.** snDNA-seq library nuclei yield comparison of fresh vs frozen nuclei of the same samples. **d-e.** Venn diagram comparing the sets of high-quality variants (**Methods)** identified in snDNA-seq libraries generated by fresh vs frozen (3 weeks) nuclei of sample PA04-2 **(e**); fresh vs frozen (14 weeks) nuclei of sample PA04-3 **(f**). The mean read depths of each library are labeled. **f.** Pseudobulk VAF comparison of all 212 shared variants between libraries generated with fresh vs frozen (14 weeks) nuclei of sample PA04-3. Key drivers preidentified by bulk WES in this case are highlighted; regression line with 90% confidence interval is drawn. **g.** Single-cell genotype heatmap of snDNA-seq libraries generated by unsorted vs sorted nuclei of sample PA04-3. The nuclei are sorted based on *KRAS* variant’s variant allele frequency (VAF) in ascending order from left to right.

**Supplementary figure 2:**
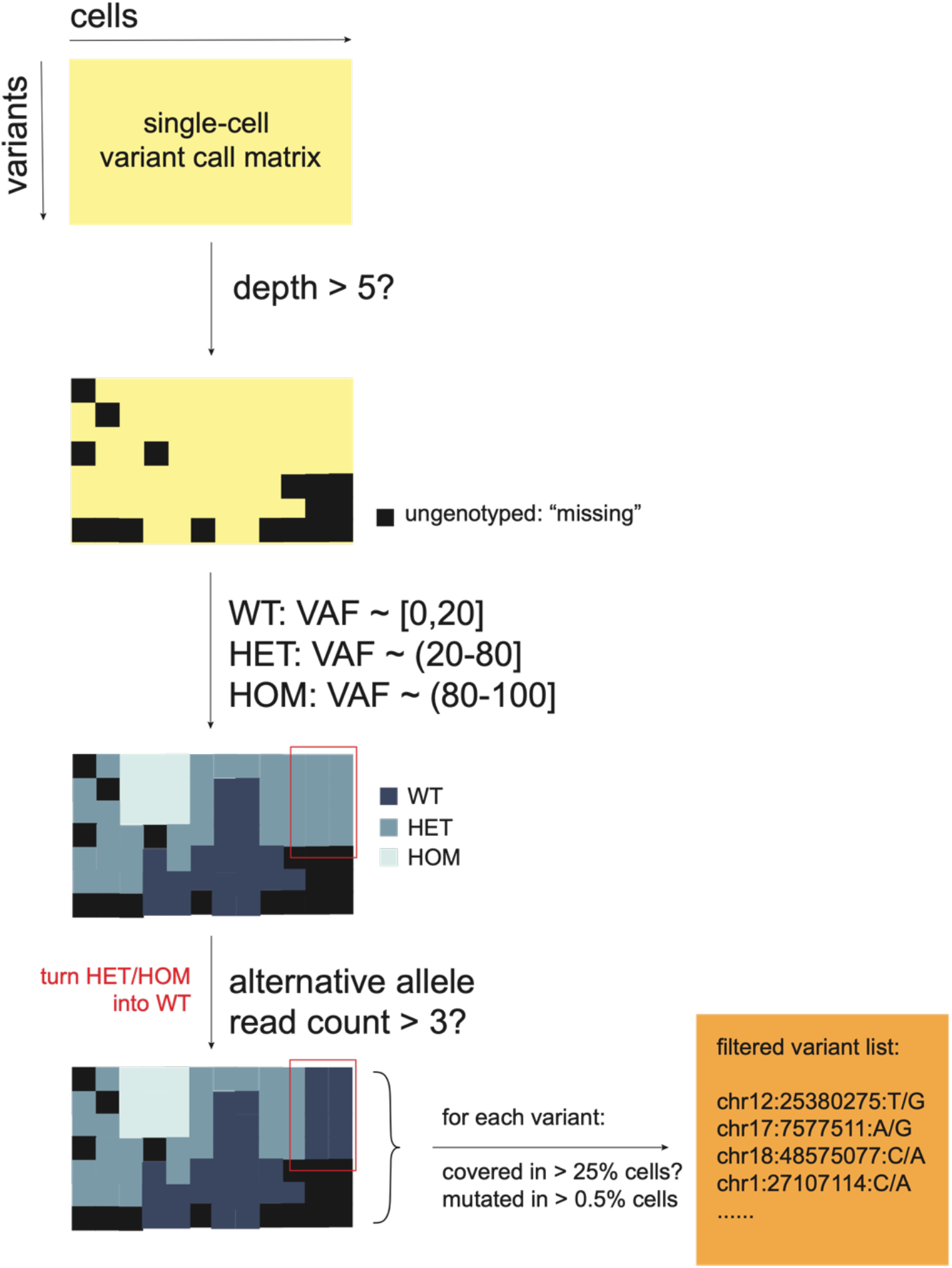
single-cell variant matrix genotyping and filtering procedure. To enable comparison of variant call sets across technical replicates, the procedure outlined in the figure and detailed in the method section was used. Briefly, each variant in each single cell was first genotyped with read depth and variant allele frequency (VAF) thresholds, and then hard filtered based on coverage and mutational prevalence to result in a final call set for each sample.

**Supplementary figure 3:**
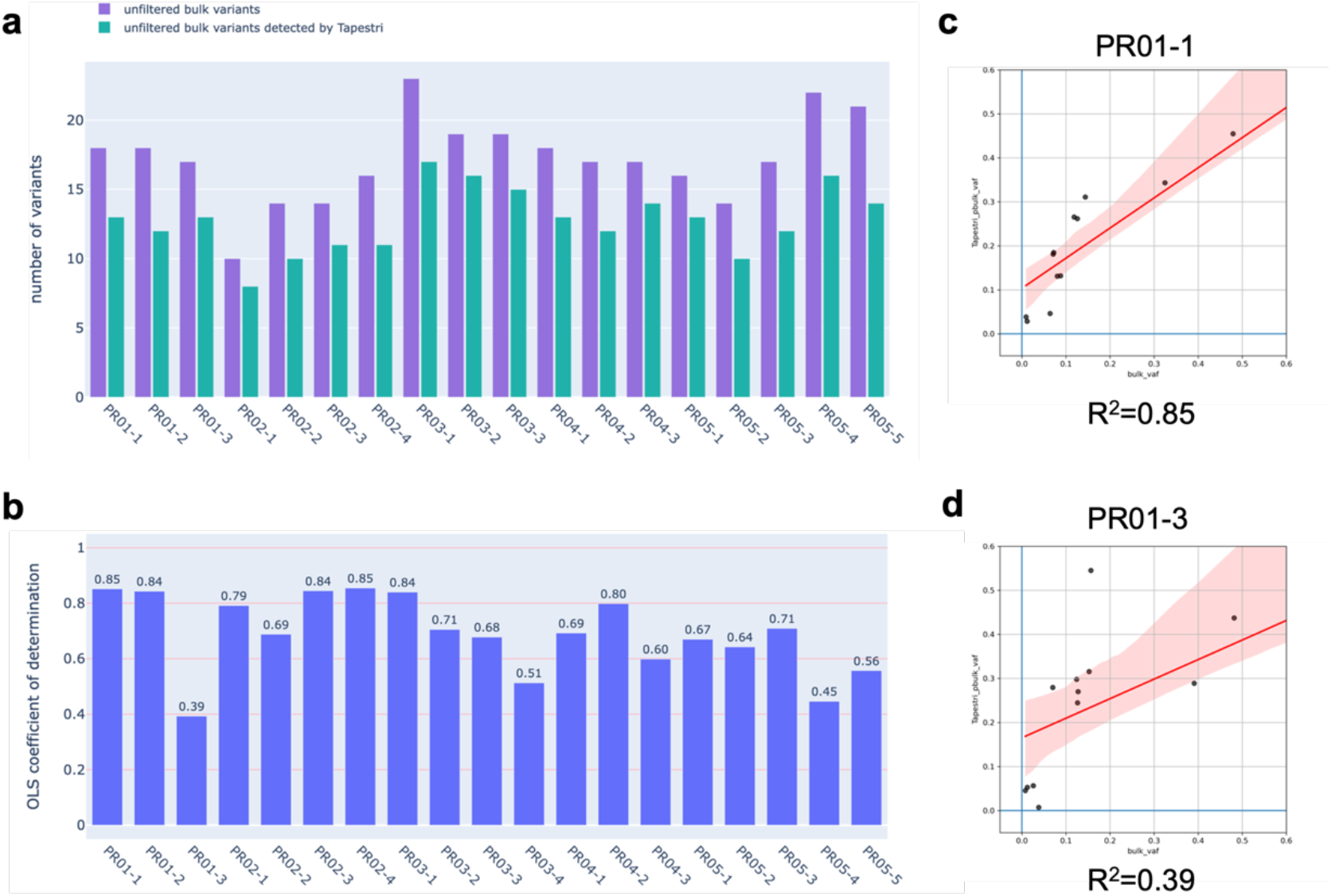
Bulk vs snDNA-seq variant comparison. **a.** For 18 samples of the same bulk WES sequencing cohort, histogram showing the number of unfiltered **(Methods)** variants called by bulk (purple) and among them, the number detected by snDNA-seq (green). **b.** Persample linear regression results of bulk VAF vs snDNA-seq pseudobulk VAF for all shared variants, except for those that have less than or equal to 2 alternative reads in bulk results; coefficient of determination (R^2^) is plotted on the y axis. **c-d.** Representative linear correlation of bulk VAF vs snDNA-seq pseudobulk VAF for shared variants for sample PR01-1 (R^2^ = 0.85), PR01-3 (R^2^ = 0.39).

**Supplementary figure 4:**
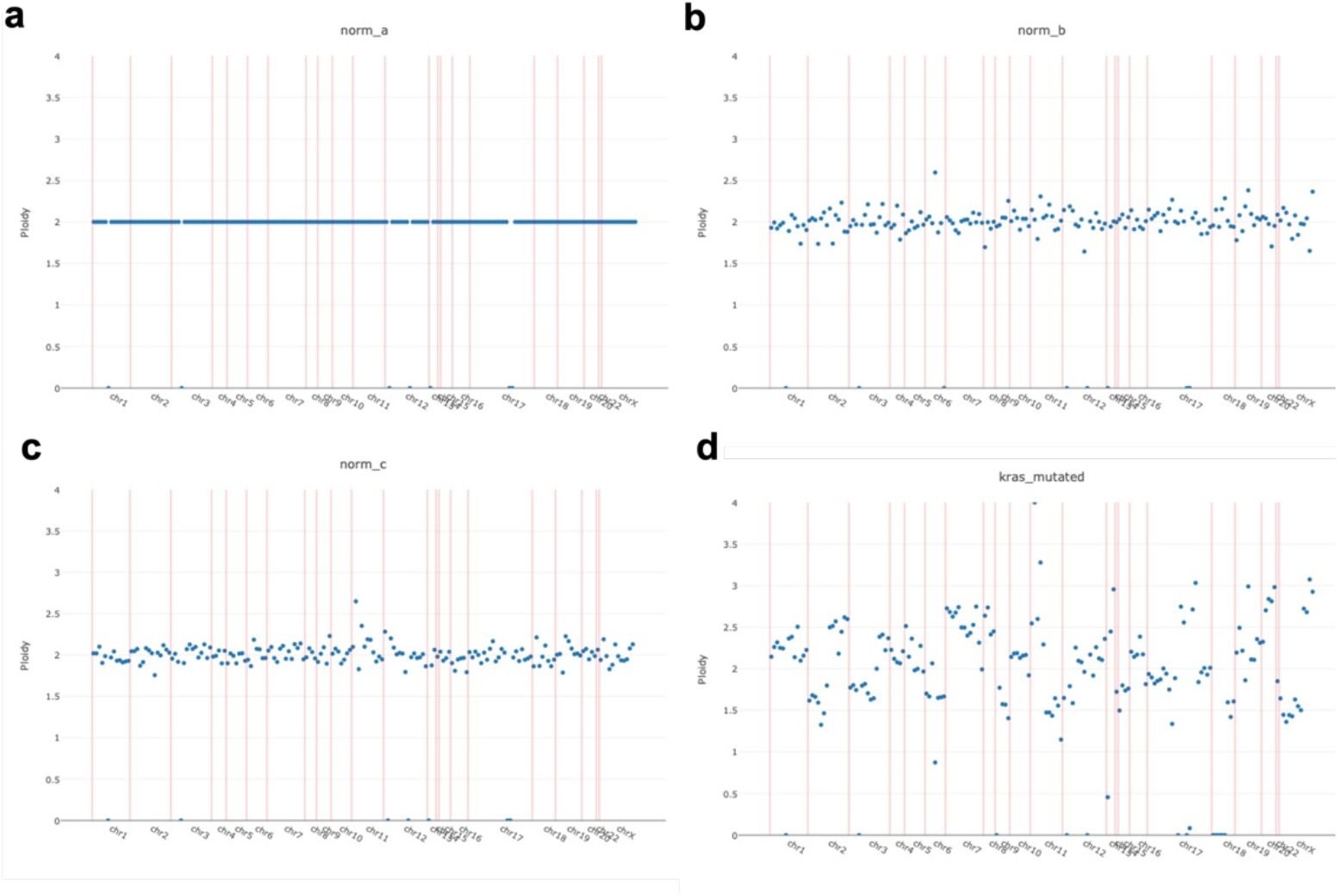
Clone-median per-amplicon ploidy for sample PA02-1. Ploidy was calculated as follows:

1. Read count per (cell * amplicon) is normalized both across cells and amplicons as described in **Methods.**
2. *KRAS*-mutated group is identified as cells carrying *KRAS* HOM/HET genotype; all other cells are assigned as putative normal cells.
3. Putative normal cells are randomly split into 3 equally-sized groups norm_a, norm_b, norm_c. Norm_a is used as diploid baseline and all other groups’ absolute ploidy is calculated as the ratio of their normalized read counts to norm_a’ normalized read counts. For sample PA02-1, the resulting absolute per-amplicon ploidy for each group is plotted above **(a-d).**

**Supplementary figure 5:**
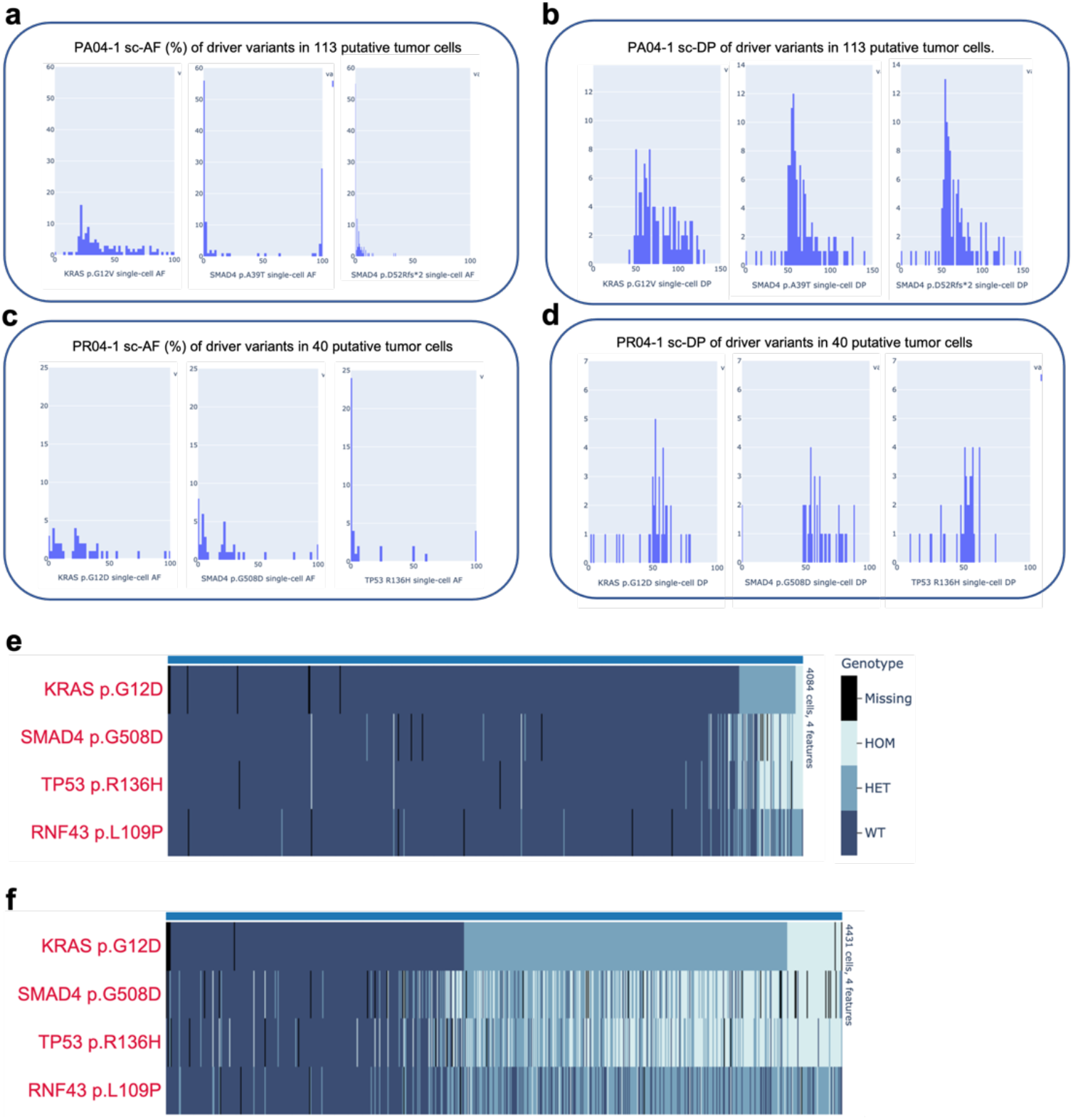
Tapestri snDNA-seq’s performance in limiting settings. **a-b.** Distribution of single-cell allele frequency (AF, **a**), read depth (DP, **b**) of the 3 main driver variants of sample PA04-1 in the 113 putative tumor cells out of 479 total cells. **c-d.** Distribution of single-cell AF (**c**), DP(**d**) of the 3 main driver variants of sample PR04-1 in the 40 putative tumor cells. **e-f.** single-cell genotype heatmap of samples PR04-2 **(e)**, PR04-3 **(f)**, which are two other samples from the same tumor as PR04-1. Cells are again sorted based on *KRAS* VAF in ascending order from left to right.

**Supplementary Figure 6:**
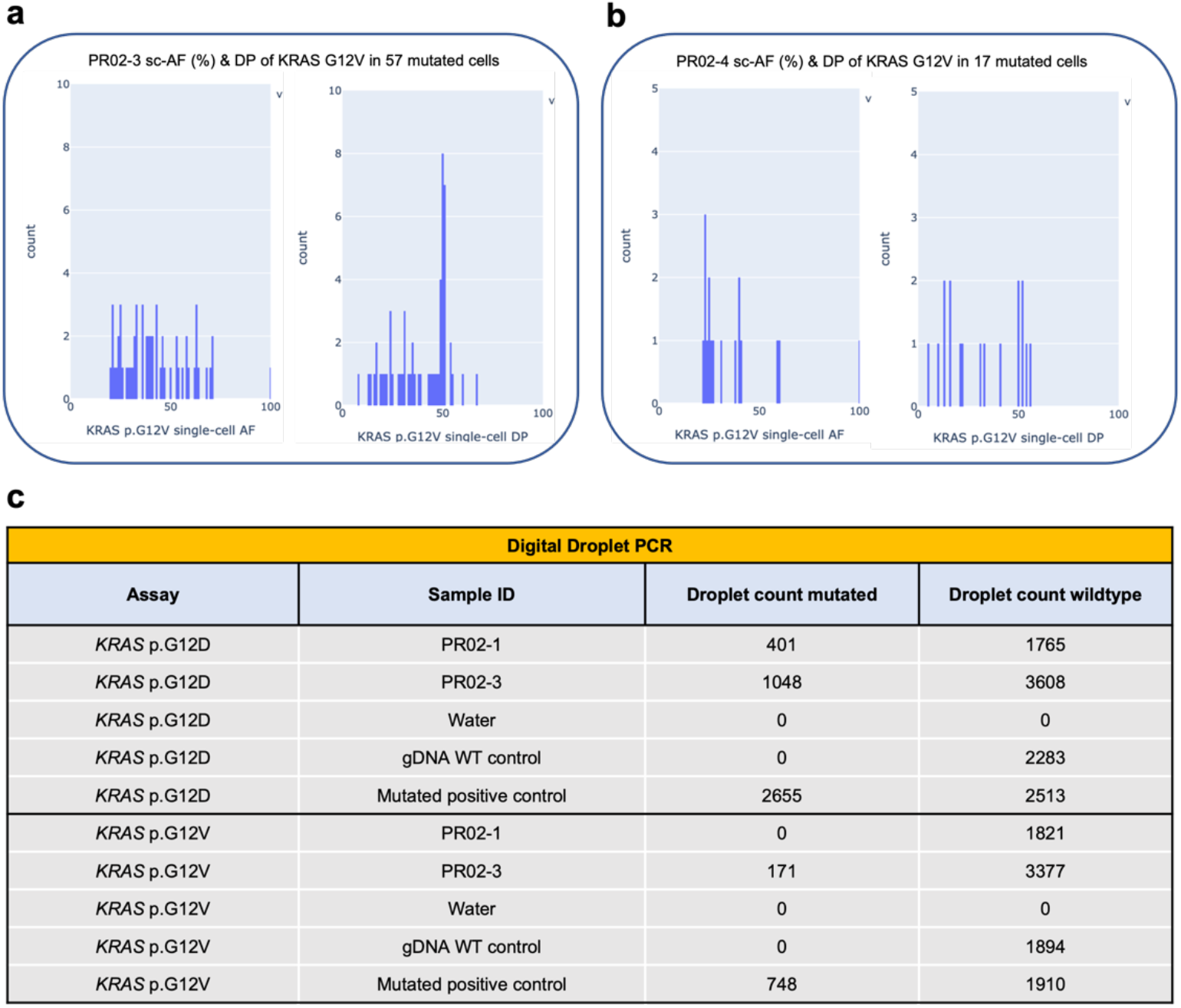
snDNA-seq identified a minor *KRAS* p.G12V mutation mutually exclusive with a major *KRAS* p.G12D mutation in the same tumor. **a-b.** Distribution of single-cell allele frequency (AF, left), read depth (DP, right) of *KRAS* p.G12V mutation in cells where it was mutated in sample PR02-3 (**a**), Pr02-4 (**b**). **c.** *D*igital droplet PCR results on *KRAS* variants in samples PR02-1 (used instead of PR02-4 because the latter’s nuclei material was depleted) and PR02-3’s leftover nuclei from Tapestri runs.

**Supplementary Figure 7:**
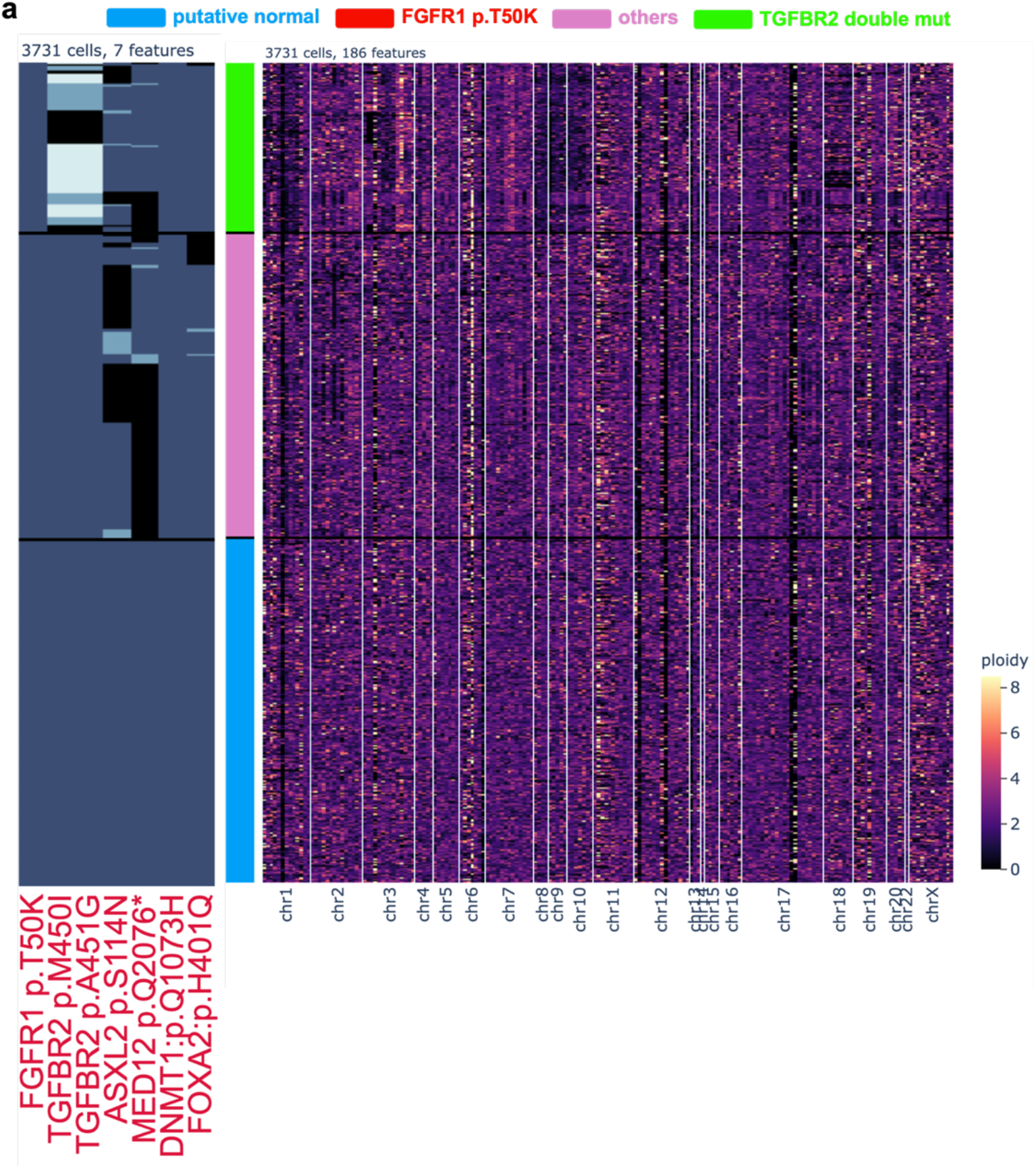
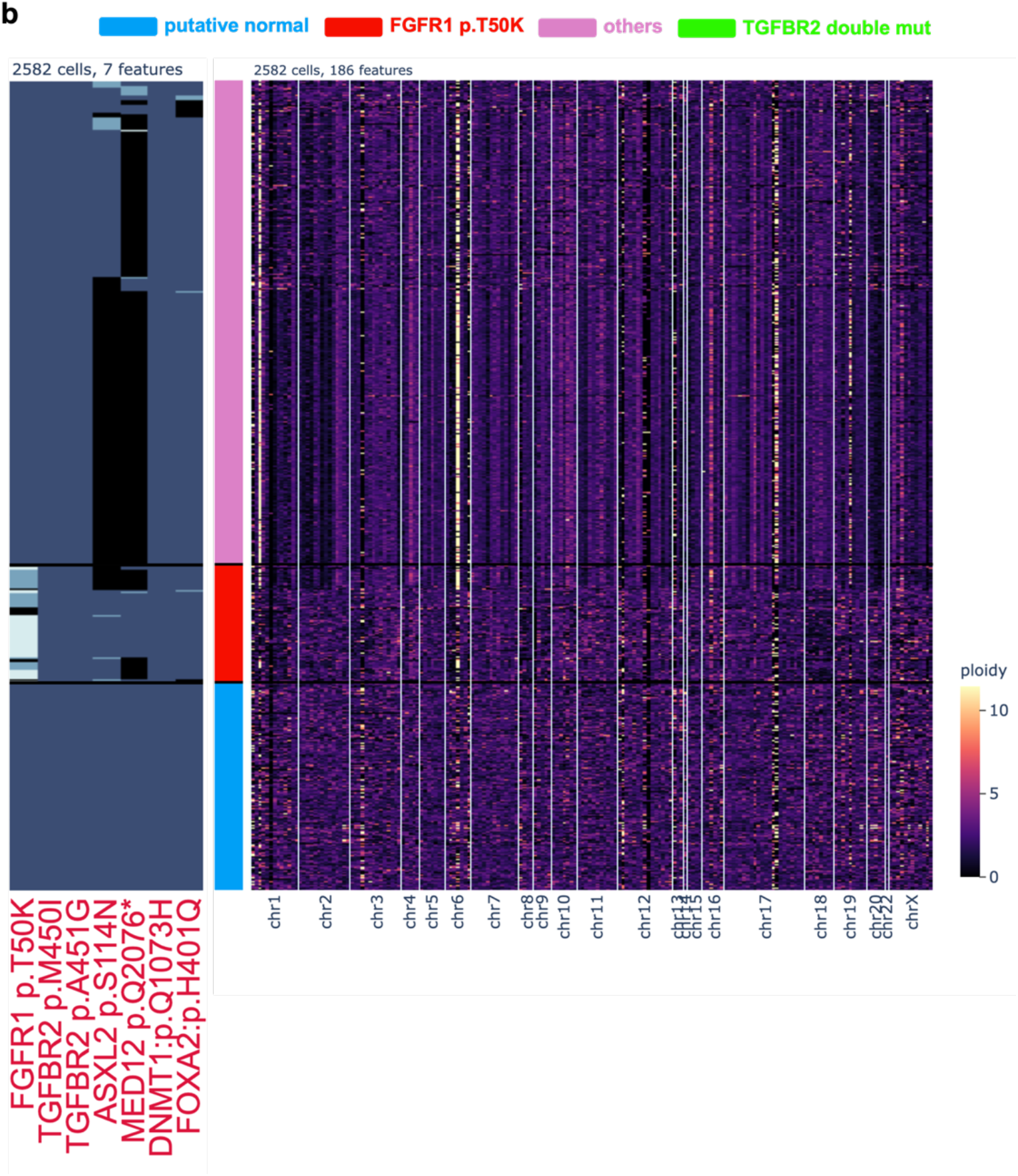
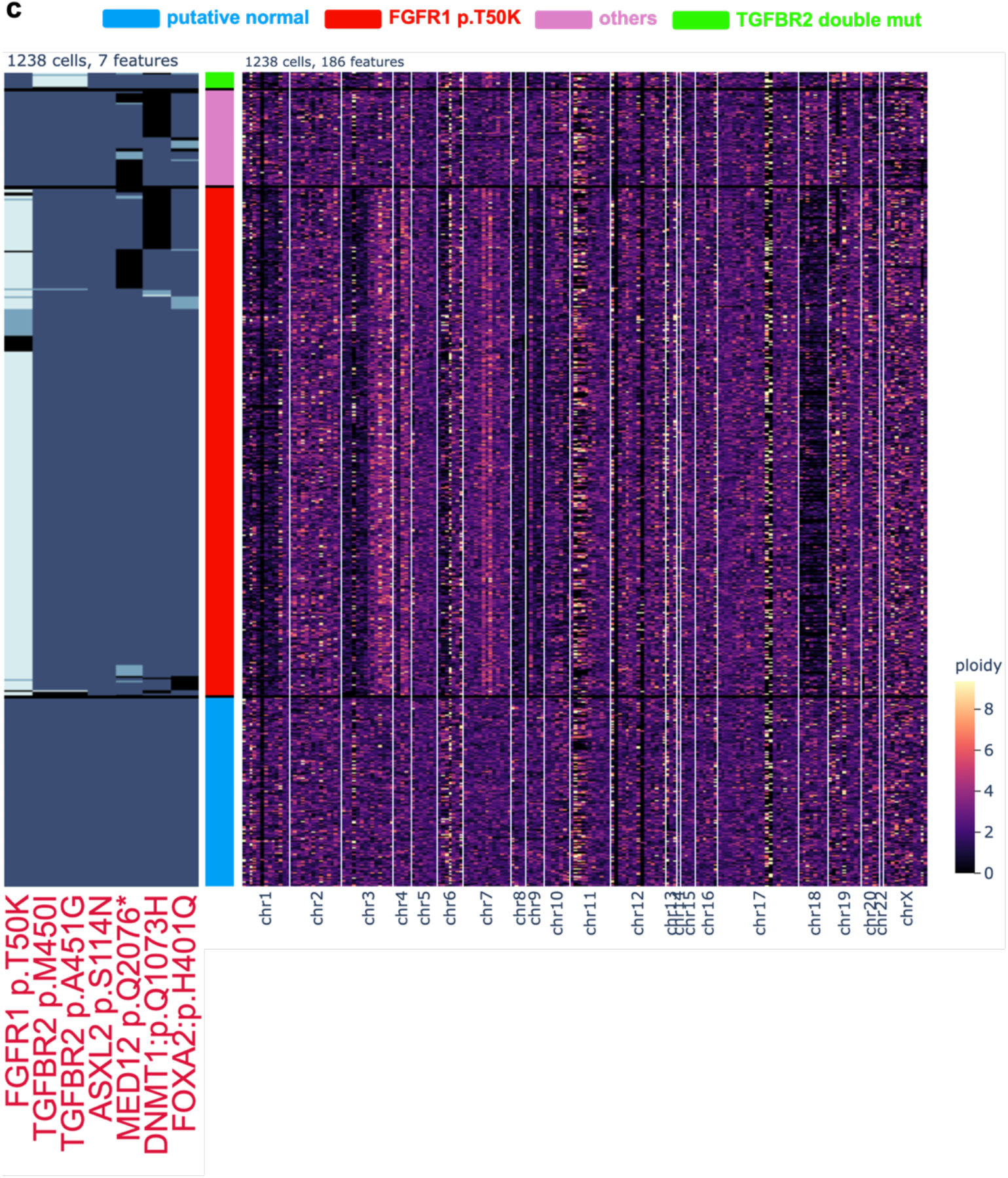
single-cell SNV and CNV results of a *KRAS* WT PDAC. **a-c.** Single-cell genotype heatmap of 7 important genomic variants pre-identified by bulk WES (left) and genomewide per-amplicon ploidy heatmap (right) for samples PR01-1 (**a**), PR01-2 (**b**), PR01-3 (**c**). Each cell’s clone identity (middle) is colored as shown by labels. The FGFR1 and TGFBR2 SNV clones are defined as cells with non-WT genotype of each gene. The “putative normal” clone is defined as cells with “WT” genotype of all 7 genetic variants. Cells are hierarchically clustered within each clone.

**Supplementary Figure 8:**
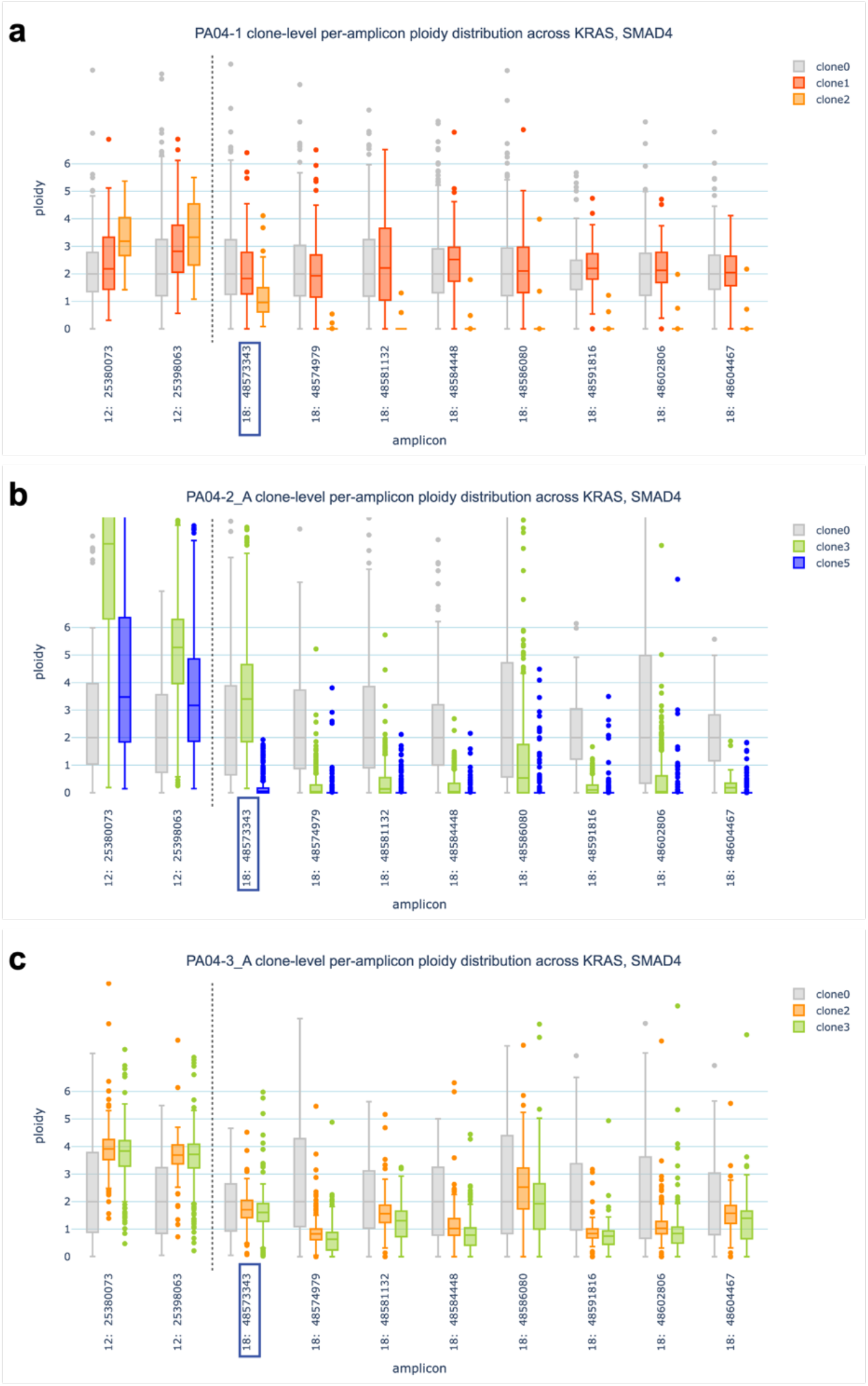
snDNA-seq revealed step-wise evolution during PDAC metastasis. **a-c.** Boxplot showing distribution of single cell ploidy for *KRAS*, *SMAD4* in sample PA04-1 (**a**), PA04-2_A (**b**), PA04-3_A (**c**), split by clone identities defined as in **Figure 6**. Each clone is colored the same as defined in **Figure 6.** Each box spans from quartile 1 (Q1) to quartile 3 (Q3). The second quartile (Q2) is marked by a line inside the box. The whiskers correspond to the box’ edges +/- 1.5 times the interquartile range (IQR: Q3-Q1). Only sample points lying outside the whiskers are shown. Sample points with ploidy > 10 are clipped from the images. Each amplicon’s starting genomic location is labeled on the x-axis and a dashed line is drawn to separate the two different genes. The amplicon where the *SMAD4* p.A39T and p.D52Rfs*2 mutations took place is outlined in blue.

