## Supplementary_Document-Doublet_estimation for "Application of high-throughput, high-depth, targeted single-nucleus DNA sequencing in pancreatic cancer"

### Doublet rate estimation

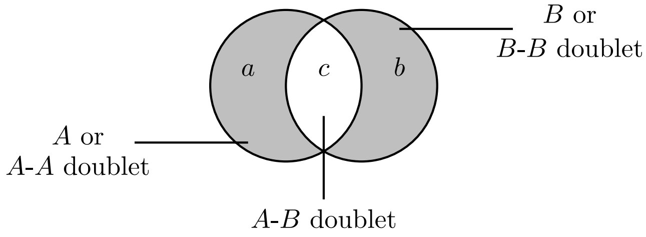


Figure 1: Venn diagram showing the different possible observations and illustrating that it is not possible to distinguish between singlets and self-doublets.

Doublets occur when multiple cells get encapsulated in the same drop and acquire the same barcode. In microfluidic experiments the doublet rate $\delta$, expected proportion of doublets in the data, depends on the rate at which the cells flow through the nozzle and time-interval in which a droplet is formed. When these quantities are small, the probability a doublet occurring can be approximated by a Binomial distribution [1–3] with a success probability $\delta$.

We start with two cell lines $A$ and $B$ with unknown proportions $x$ and $y$, respectively. Since there are only two cell lines in the mixture, we have $x + y = 1$. Let $n$ be the total number of observations, out of which $a$ contain mutations from cell line $A$, $b$ contain mutation from cell line $B$ and $c$ contain mutations from both cell line $A$ and $B$ (see Fig. 1). As such, $a + b + c = n$. Our goal is to compute the doublet rate $\delta$ for given values of $a$, $b$ and $c$.

We assume that the number of observation is much smaller than the size of the initial mixture from which the cells are sampled. The expected total number of singlets is $n(1 - \delta)$, with number of singlets of cell line $A$ is $x(1 - \delta)n$ and the number of singlets of cell line $B$ is $y(1 - \delta)n$. Moreover, we assume that each cell in the doublet is picked independently from the mixture. A doublet can either be a neotypic $A$-$B$ doublet, comprising of one cell from cell line $A$ and one cell from cell line $B$, or a self-doublet $A$-$A$ or $B$-$B$ comprising of two cells from the cell line $A$ or $B$, respectively. The expected number of $A$-$A$ doublets is $x^2\delta n$, the expected number of $B$-$B$ doublets is $y^2\delta n$ and the expected number of $A$-$B$ doublets is $2xy\delta n$. Note that the expected total number of doublets is $n\delta$. We need to solve the following set of nonlinear equations

$$\begin{aligned} xn(1 - \delta) + x^2\delta n &= a \\ yn(1 - \delta) + y^2\delta n &= b \\ 2xy\delta n &= c, \\ a + b + c &= n. \end{aligned}$$

We solve these nonlinear equations numerically using the open-source package SCIPY.

#### References

[1] Hamim Zafar, Anthony Tzen, Nicholas Navin, Ken Chen, and Luay Nakhleh. Sifit: inferring tumor trees from single-cell sequencing data under finite-sites models. *Genome biology*, 18(1):

1–20, 2017.

[2] Samuel L Wolock, Romain Lopez, and Allon M Klein. Scrublet: computational identification of cell doublets in single-cell transcriptomic data. *Cell systems*, 8(4):281–291, 2019.

[3] Leah L Weber, Palash Sashittal, and Mohammed El-Kebir. doubletd: detecting doublets in single-cell dna sequencing data. *Bioinformatics*, 37(Supplement_1):i214–i221, 2021.
